# Validation of MHPC512: A Publicly Available Special-Purpose Supercomputer for Molecular Dynamics Simulations of Biomolecular Systems

**DOI:** 10.64898/2026.06.10.727820

**Authors:** Lin Chen, Yuhan Chen, Ziqi Cheng, Jing Guo, Ming He, Huan Li, Xinyu Li, Zhen Li, Jule Ma, Shuo Ma, Can Peng, Cheng Qian, Zhi Qu, Xian-gang Sun, Xuan Tang, Yunfei Wang, Bing Yu, Yuzhuo Zhai, Bingyao Zhang, Shang Zhang, Shuo Zhang, Zhonghan Hu, Yibing Shan, Ye Mei

## Abstract

MHPC512 is a massively parallel, special-purpose supercomputer designed primarily for atomic-level molecular dynamics (MD) simulations of biomolecular systems. It comprises 512 processor units interconnected by a high-speed three-dimensional torus network and employs a custom chip architecture that uses 35-bit fixed-point arithmetic to accelerate computation while controlling precision loss within an acceptable margin. The preprocessor is compatible with GROMACS and AMBER input formats and supports widely used biomolecular force fields (including CHARMM, AMBER, and OPLS/AA), the Neutral Territory method for short-range nonbonded interactions, the k-space Gaussian Split Ewald method for long-range electrostatics, and multiple thermostats, barostats, and integrators. We present a three-tier validation protocol—comparing static energy and virial components, examining ensemble distributions (NVE, NVT, NPT), and evaluating long-time statistical properties—demonstrating that MHPC512 reproduces results consistent with GROMACS and AmberTools. Application examples, including bulk water, dipeptide conformational sampling, folding of fast-folding peptides, membrane–protein systems, lipid self-assembly, and GPCR conformational transitions, further confirm its reliability. MHPC512 has been deployed at multiple supercomputing centers and is publicly accessible, representing a significant advance in high-throughput, large-scale biomolecular MD simulations.

## 1 Introduction

Molecular dynamics (MD) simulation has become an indispensable tool across a wide range of scientific disciplines, with particularly extensive applications in biology, biopharmaceuticals, and materials science.^1–4^ By numerically solving Newton’s equations of motion, MD enables researchers to explore the behavior of atomic systems with exceptional detail—both spatially and temporally—providing insights into phenomena that are often difficult or impossible to observe experimentally. Its sub-nanometer spatial resolution and picosecond temporal resolution allow for the detailed examination of molecular interactions and dynamic processes, making MD a powerful complement to experimental techniques. In this way, MD facilitates hypothesis generation, mechanism elucidation, and the validation of experimental results.

Despite these advantages, MD simulations face inherent limitations. ^5–7^ While offering unparalleled resolution, the accessible timescales and length scales remain orders of magnitude smaller than those of many biologically or chemically relevant processes. For example, conformational changes in proteins may occur on the millisecond timescale, and drug unbinding from a target site may take minutes or even hours—far beyond the reach of current MD simulations on conventional supercomputers. Bridging this scale gap has been a long-standing challenge and a central focus of computational research.^8–10^

Over the past two decades, significant progress has been made in general-purpose graphics processing units (GPGPUs) and their associated software, making GPU-accelerated MD a standard solution. However, speedup remains limited.^11^ To overcome this barrier, researchers have explored a variety of hardware and algorithmic innovations. Early efforts involved the use of Field-Programmable Gate Arrays (FPGAs) to accelerate specific calculations. The GRAPE project pioneered dedicated hardware solutions for MD by implementing custom very-large-scale integration (VLSI) chips and later FPGAs, demonstrating the potential of purpose-built accelerators for molecular dynamics.^12,13^ Subsequent developments have significantly advanced the field through more efficient algorithms and the use of specialized hardware. A prominent example is the Anton series of supercomputers developed by D. E. Shaw Research (DESRES), which are purpose-built for MD simulations. Anton 3, the latest in the series, achieves an order-of-magnitude improvement in performance over Anton 2 and is over 100 times faster than the available general-purpose systems when Anton 3 was released.^14–16^

Anton 3 incorporates several architectural innovations, including a novel compression algorithm that predicts atomic positions based on past trajectories, greatly enhancing bandwidth efficiency. This is complemented by improvements in the off-chip communication network and hardware accelerators optimized for pairwise atomic interactions. As a result, Anton 3 (512 nodes) can routinely simulate million-atom systems at rates exceeding 100 *µ*s/day, enabling the investigation of biologically relevant events that require multi-microsecond trajectories. Additional performance gains are achieved through specialized Pairwise Point Interaction Pipelines (PPIPs), tailored for varying interaction distances to maximize computational throughput. Unfortunately, Anton 3 is almost exclusively used in DESRES and is not accessible to the wider research community, with the sole exception of Pittsburgh Supercomputing Center.

While Anton 3 represents the current peak in specialized hardware for MD simulations, it is not the only system designed with this purpose in mind. In this manuscript, we introduce MHPC512, a publicly available, massively parallel, special-purpose supercomputer designed for molecular dynamics simulations. We begin by describing the molecular dynamics algorithms available on MHPC512, followed by validation studies that demonstrate its accuracy and consistency with established MD software. Finally, we present several application examples to showcase the system’s performance and capabilities.

## 2 Technique Details

MHPC512 consists of 512 processor units^17^, arranged in a three-dimensional (3D) topology connected via a high-speed 3D-torus network. Each processor unit contains 36 cores, and atoms in the simulation box are spatially mapped to these cores based on their fractional coordinates in the simulation box. To accelerate arithmetic operations, fixed-point number representations are adopted in place of floating-point arithmetic. This engineering effort achieves comprehensive performance optimization across four dimensions: data structure, read/write caching, redundancy elimination, and distributed parallelism. While maintaining strict compliance with accuracy requirements, it substantially enhances overall computational efficiency, reduces communication overhead, and ensures efficient utilization of both computing power and bandwidth resources.

The original 64-bit double-precision floating-point operations are iteratively transformed into 35-bit fixed-point arithmetic, except for the force components, which are calculated and transferred in 70-bit format. Within a controlled and acceptable precision loss margin, this optimization markedly improves computational performance. Moreover, data transmission employing the 35-bit fixed-point format significantly reduces the volume of data transferred, thereby jointly optimizing arithmetic throughput and transmission efficiency.

Through hierarchical storage and optimized read/write mechanisms, data throughput is enhanced. Intermediate computational variables are uniformly stored in core registers, substantially reducing performance penalties caused by frequent memory accesses. A multi-level cache management scheme is adopted: high-frequency data resides in high-speed cache, whereas low-frequency data is placed in L3 cache, aligning storage hierarchy with data access patterns. Additionally, scattered atomic read/write operations are consolidated into batch transactions, effectively improving overall read/write bandwidth utilization.

Ineffective computations are streamlined via four strategies: preprocessing, hardware acceleration, precise domain control, and algorithmic iteration. All bonded interactions are preprocessed and allocated to specific chips in advance, thereby avoiding repeated retrieval of bonded interactions with target chips and reducing redundant calculations. For long- and short-range nonbonded interactions, the conventional full-neighbor-list traversal is discarded. Instead, a dedicated transmission structure is designed, which, together with a self-developed co-processor, performs hardware-based filtering—completely circumventing inefficient global traversal and significantly boosting computational efficiency. Furthermore, transmission ranges are precisely delimited according to system scale. Based on the box length and non-bonded truncation distance, the required transmission blocks are carefully designed for atomic systems of 300k, 500k, 800k, and 1 million atoms, achieving lower data transfer volumes in large-scale systems. Finally, a highly efficient, simple high-speed parallel logic replaces traditional inefficient conditional branching, further accelerating the computational pipeline.

Maximal utilization of parallel computing power is achieved through heterogeneous task load balancing, differentiated mode selection, and lightweight data management. First, computational tasks on heterogeneous devices are rationally partitioned and allocated. Pre-assignment of bonding computations balances computation duration across processes, completely eliminating idle waiting. Second, a scale-aware heterogeneous strategy governs the computation–transmission trade-off. For systems with less than 260k atoms, a strategy of “computation replacing transmission” is adopted, where pairwise bidirectional recomputation reduces data transmission overhead; for systems with 260k atoms and above, a strategy of “transmission replacing computation” is employed, where a small amount of force back-transmission overhead is traded for a significant reduction in computational load. Finally, data storage and migration policy is optimized. Each process holds only its own domain-specific particle information, abandoning the full-system data copying paradigm. This achieves on-demand migration of computational data without redundant backups, effectively reducing memory occupancy and data redundancy overhead.

MHPC512 is compatible with input file formats from GROMACS^18^ and AMBER^19,20^ for atomic positions, velocities, cell dimensions, interaction parameters, and topologies. It provides full support for widely used biomolecular force fields, including CHARMM^21^, AMBER^22^, and OPLS/AA^23^. For short-ranged nonbonded interactions—such as truncated Lennard-Jones potentials and real-space electrostatics—MHPC512 implements the Neutral Territory (NT) method^24,25^. This approach significantly reduces inter-processor communication by allowing pairwise force calculations between particles located on different nodes to be assigned to a third, neutral node—rather than computing them exclusively on one of the two home nodes, as in traditional spatial decomposition schemes. Long-range electrostatic interactions are handled using the *k*-space Gaussian Split Ewald (*k*-GSE) method, originally developed by Shan et al. for Anton.^26^ In this approach, the electrostatic potential is decomposed into a rapidly decaying short-range term—computed directly in real space using blurred Gaussian countercharges—and a long-range term, which is calculated in reciprocal space via charges projected onto mesh grids.

MHPC512 supports both velocity-Verlet and leap-frog integrators. In addition, the reversible reference system propagator algorithm (r-RESPA)^27^ is implemented to accelerate simulations by separating the force field into fast- and slow-varying components via Trotter expansion^28^. In the simulations shown below, the slow-varying electrostatic component is updated every two integration steps. M-SHAKE^29^ and RATTLE^30^ algorithms are available to constrain bonds involving hydrogen atoms. To control temperature, both Berendsen^31^ and Nosé–Hoover chain (NHC) thermostats^32^ are implemented. For pressure regulation, MHPC512 supports Berendsen^31^ and Parrinello–Rahman^33^ barostats, with options for isotropic and semi-isotropic coupling—the latter being especially useful for membrane systems. As for isotropic pressure coupling, Andersen barostat^34^ is also available. Additionally, surface-tension coupling^35^ is available to control membrane geometry in the XY-plane. Both cubic and orthorhombic simulation boxes are supported.

While computational efficiency is essential, accuracy and reliability remain the top priorities. Before MHPC512 can be considered a trustworthy platform for scientific MD simulations, rigorous validation is required against widely used reference software and hardware. Our validation protocol comprises three steps. As the first step of validation, static tests compare the individual components of the energy and virial tensors computed by MHPC512 with those generated by GROMACS v2024.3 and AmberTools24^36^, using identical input structures, parameters, and topologies. Directional decomposition of the virial tensor is also assessed, which is critical for semi-isotropic pressure coupling. Although correct energy and force are necessary for a reliable simulation, they are not enough since integrators, thermostat and barostat methods are also critical, especially for long time simulations since more than 10^9^ steps of propagation are normally required and the cumulative error may deteriorate the simulation. Therefore, in the second step, the energy conservation in constant number of particles, volume and energy (NVE) simulations, energy distribution with respect to different temperatures in constant number of particles, volume and temperature (NVT) simulations, and volume distributions as a function of external pressure in constant number of particles, pressure and temperature (NPT) simulations are examined. In the third step, extended MD simulations are conducted to evaluate statistical properties across ensembles, which include structural properties of bulk water, Ramachandran plots of some dipeptides, folding simulations of some short peptides, and structures of membrane and membrane proteins. We believe this three-tier validation process comprehensively covers the most common use cases of conventional MD simulations. Finally, to demonstrate the practical utility of MHPC512, we present results from real-world applications that illustrate its ability to tackle challenging scientific problems. In the following, only the simulation results are presented, with simulation setups detailed in the supporting information.

## 3 Results and Discussion

### 3.1 Potential Energy and Virial

This study evaluated a broad array of force fields beyond OPLS/AA^23,37,38^, encompassing both CHARMM and AMBER families. Specifically, force fields tested include CHARMM22*^39^, CHARMM27^40^, CHARMM36^41^, CHARMM36m^42^, AMBER99SB-ILDN,^43^ AMBER14SB^44^, AMBER19SB^45^, GAFF2^46^ for organic small molecules, LIPID17 and LIPID21^47^ for lipids, Parmbsc1^48^, OL15^49^, OL21^50^, OL24^51^, Tumuc1 DNA^52^, OL3^53,54^, and RNA.Shaw^55^ for nucleic acids, GLYCAM 06j-1^56^ for carbohydrates, and the phosphorylated-residue extensions phosaa10^57^, phosaa14SB^58^, and phosaa19SB^58^. For water models, TIP3P (original and CHARMM mTIP3P)^59,60^, SPC^61,62^, SPC/E^63^, OPC3^64^ and SPC/Fw^65^ were assessed. MHPC512 supports input files in both AMBER and GROMACS formats, allowing users to migrate seamlessly from either platform.

Tables 1 and 2 compare MHPC512 with GROMACS for the potential energy and virial of Janus kinase 2, and with AmberTools for the potential energy of a POPC lipid bilayer, respectively. Both systems comprise approximately 100,000 atoms. Input files follow the conventions of the reference programs (GROMACS for Janus kinase 2 and AMBER for the lipid bilayer), and MHPC512 can read either format directly. The relative error in potential energy is negligible across all cases. The virial reported by MHPC512 is 2.0000 times of GROMACS, because their definitions differ by a factor of 2. A comprehensive list of validated force fields is available in the Supporting Information. These results confirm that MHPC512 correctly parses force-field parameters and accurately computes the corresponding interactions.

**Table 1:**
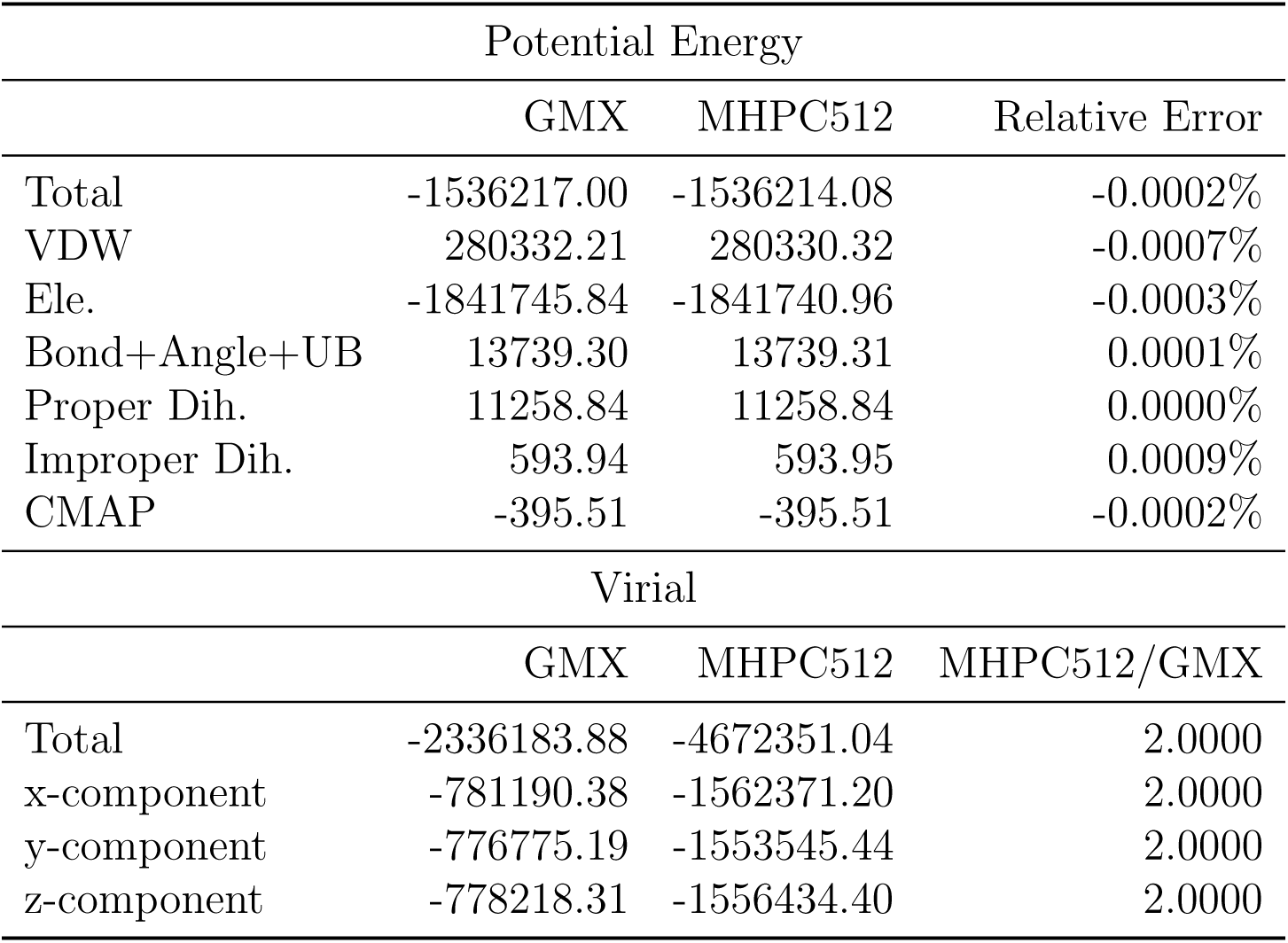
Potential energy and virial (kJ/mol) of Janus kinase 2 (PDB ID: 8C09) ^66^ parameterized with CHARMM36m force field. Input files are in the GROMACS format.

**Table 2:**
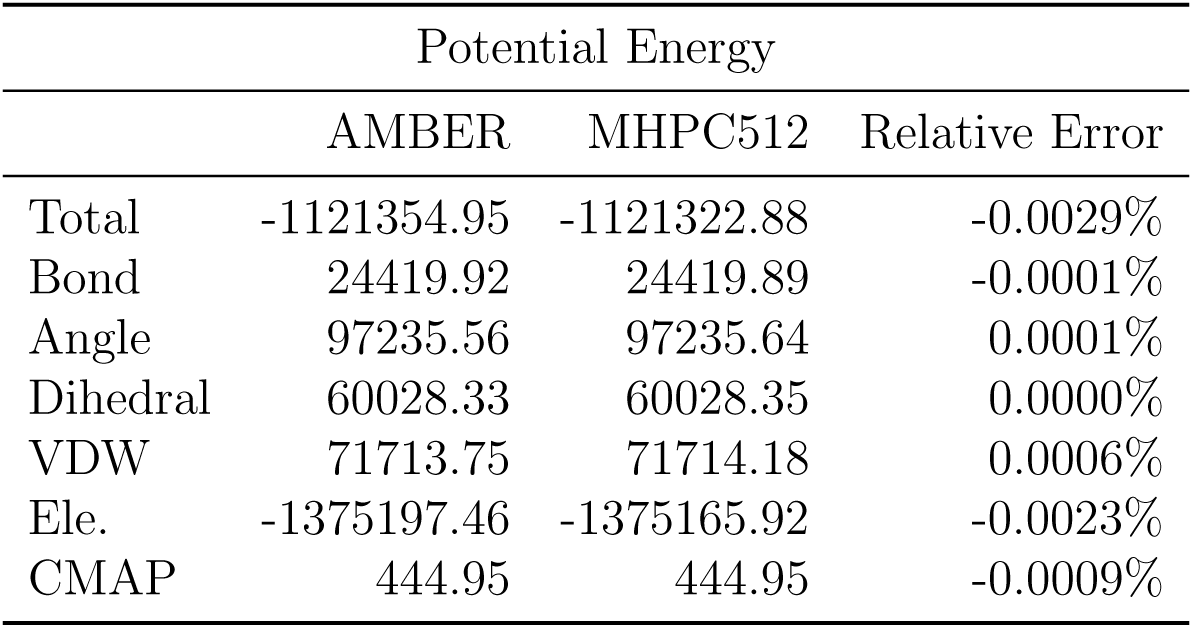
Potential energy (kJ/mol) of a POPC bilayer in water parameterized with Lipid21 and TIP3P force fields. Input files are in the AMBER format.

|  | Potential Energy |  |  |
| --- | --- | --- | --- |
|  | AMBER | MHPC512 | Relative Error |
| Total | -1121354.95 | -1121322.88 | -0.0029% |
| Bond | 24419.92 | 24419.89 | -0.0001% |
| Angle | 97235.56 | 97235.64 | 0.0001% |
| Dihedral | 60028.33 | 60028.35 | 0.0000% |
| VDW | 71713.75 | 71714.18 | 0.0006% |
| Ele. | -1375197.46 | -1375165.92 | -0.0023% |
| CMAP | 444.95 | 444.95 | -0.0009% |

### 3.2 Ensemble Tests

The NVE ensemble verification focuses on examining the energy drift rate of MHPC512 during long-time simulations and comparing it to the corresponding properties using GROMACS. In GROMACS simulations, updating the neighbor list at each step minimizes energy drift caused by van der Waals (VDW) omissions. Similarly, MHPC512 simulations were processed accordingly. As shown in Fig. 1, the rate of kinetic energy drift on MHPC512 (0.77 J*/*mol *·* ns) falls between that of mixed precision (7.07 J*/*mol *·* ns) and double precision (0.00 J*/*mol *·* ns) GROMACS. This is due to the fixed-point computation scheme used on MHPC512, which has numerical precision lower than double-precision floating-point numbers but higher than single-precision floating-point numbers. Since mixed-precision GROMACS simulations are accepted by the community, the precision of MHPC512 can also be considered acceptable.

**Figure 1:**
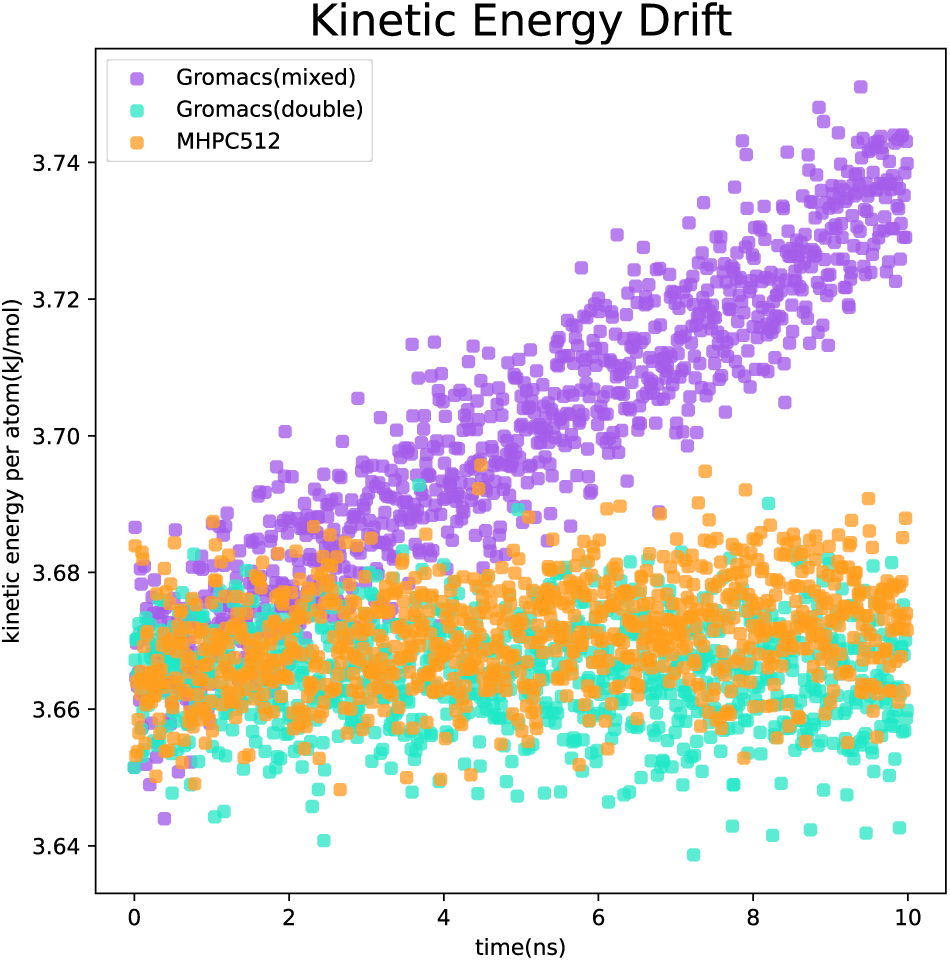
Kinetic energy drift per atom in the simulations using GROMACS (mix precision and double precision) and MHPC512.

The NVT ensemble verification focuses on examining whether the potential energy distribution obtained from long-time simulations at 298.5 K and 300.0 K on MHPC512 aligns with the given ensemble and comparing it to the corresponding results from GROMACS using similar algorithms. In the MHPC512 and GROMACS simulations, the M-SHAKE algorithm^29^ and SETTLE algorithm^67^ were applied respectively to constrain the bond lengths and angles in water molecules. As shown in Fig. 2, MHPC512 demonstrates slightly lower reliability than GROMACS in NVT simulations.

**Figure 2:**
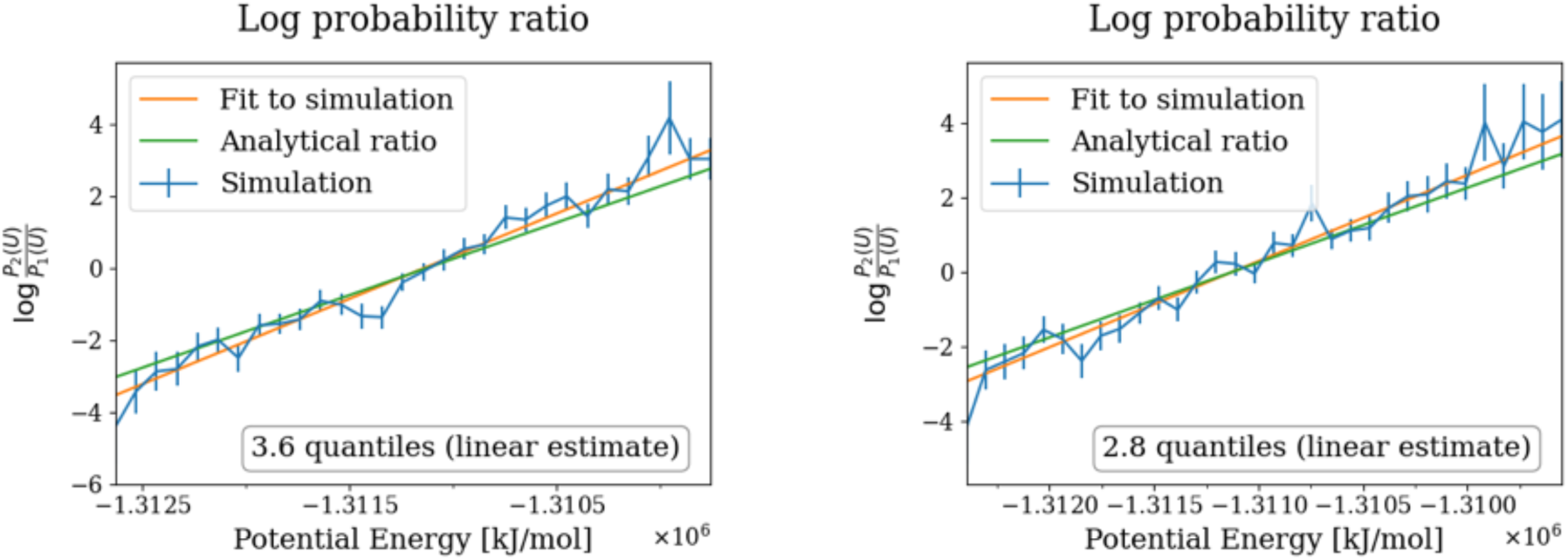
Potential energy distributions from the NVT simulations of (left) MHPC512 and (right) GROMACS on Nvidia RTX 4090D. Nosé-Hoover chain thermostat is used for temperature regulation.

The NPT ensemble verification focuses on examining whether the volume distribution obtained from long-time simulations at 1 bar and 50 bar (with temperature maintained at 300 K) on MHPC512 aligns with the given ensemble and comparing it to the corresponding results from GROMACS using similar algorithms. Bond constraints were also applied to the water molecules as in the NVT simulations. As shown in Fig. 3, MHPC512 demonstrates reliability comparable to GROMACS in NPT simulations.

**Figure 3:**
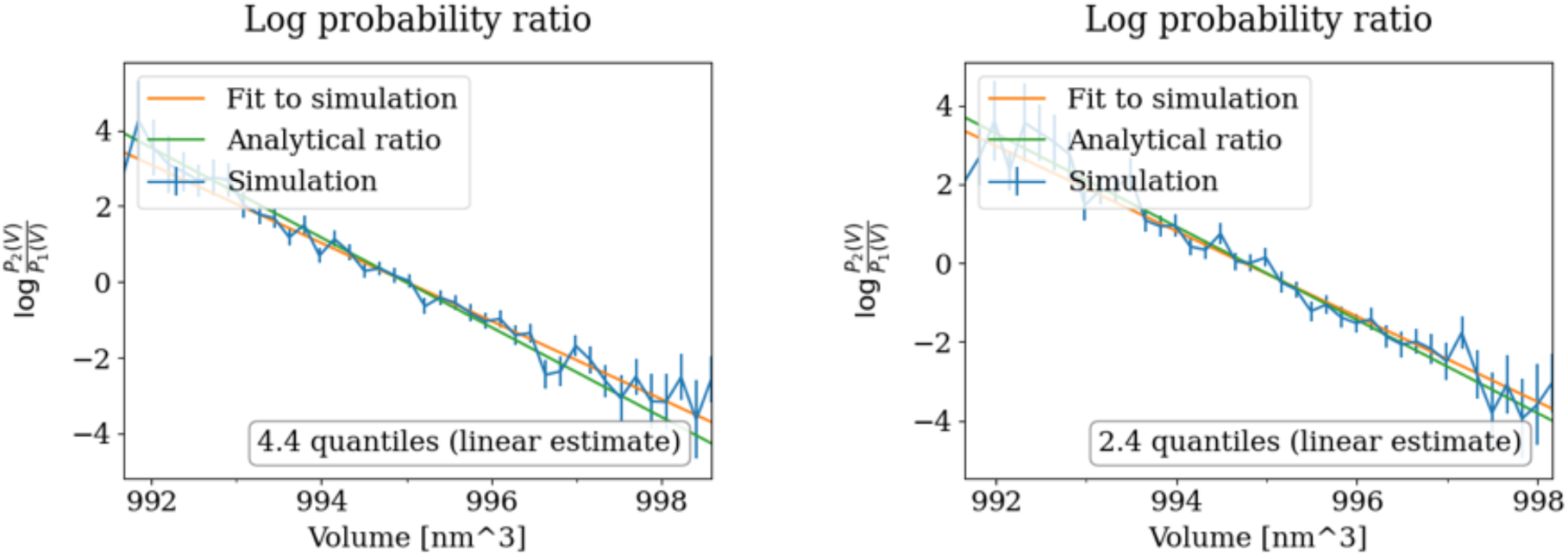
Volume distributions from the NPT simulations of (left) MHPC512 and (right) GROMACS on Nvidia RTX 4090D. Nosé-Hoover chain thermostat and Parrinello-Rahman barostat are used for temperature and pressure regulations.

### 3.3 Structure of SPC/E Water Box

The structural characteristics of an SPC/E water box were investigated after a 100-ns NPT simulation on MHPC512 and compared with those from a simulation using GROMACS. The temperature and pressure were regulated using the Berendsen algorithm. The results are presented in Fig. 4. It can be observed that both simulations exhibit similar diffusion behaviors for oxygen atoms, with only minor differences that may arise from the difference between number representations (floating point vs. fixed point) in the two platforms. Additionally, the rotational dynamics of water molecules show consistency between the two simulations. In terms of radial distribution functions, analyses based on 100-ns trajectory data (with statistics averaged every 10 ns) reveal that the peak positions of the first peak (0.18 nm) and second peak (0.32 nm) of the O-H radial distribution function are consistent. Similarly, the characteristic peak (0.28 nm) and the second layer peak (0.45 nm) of the O-O radial distribution function align well with each other. The consistency observed in these results validates the reliability of the simulation conducted on MHPC512.

**Figure 4:**
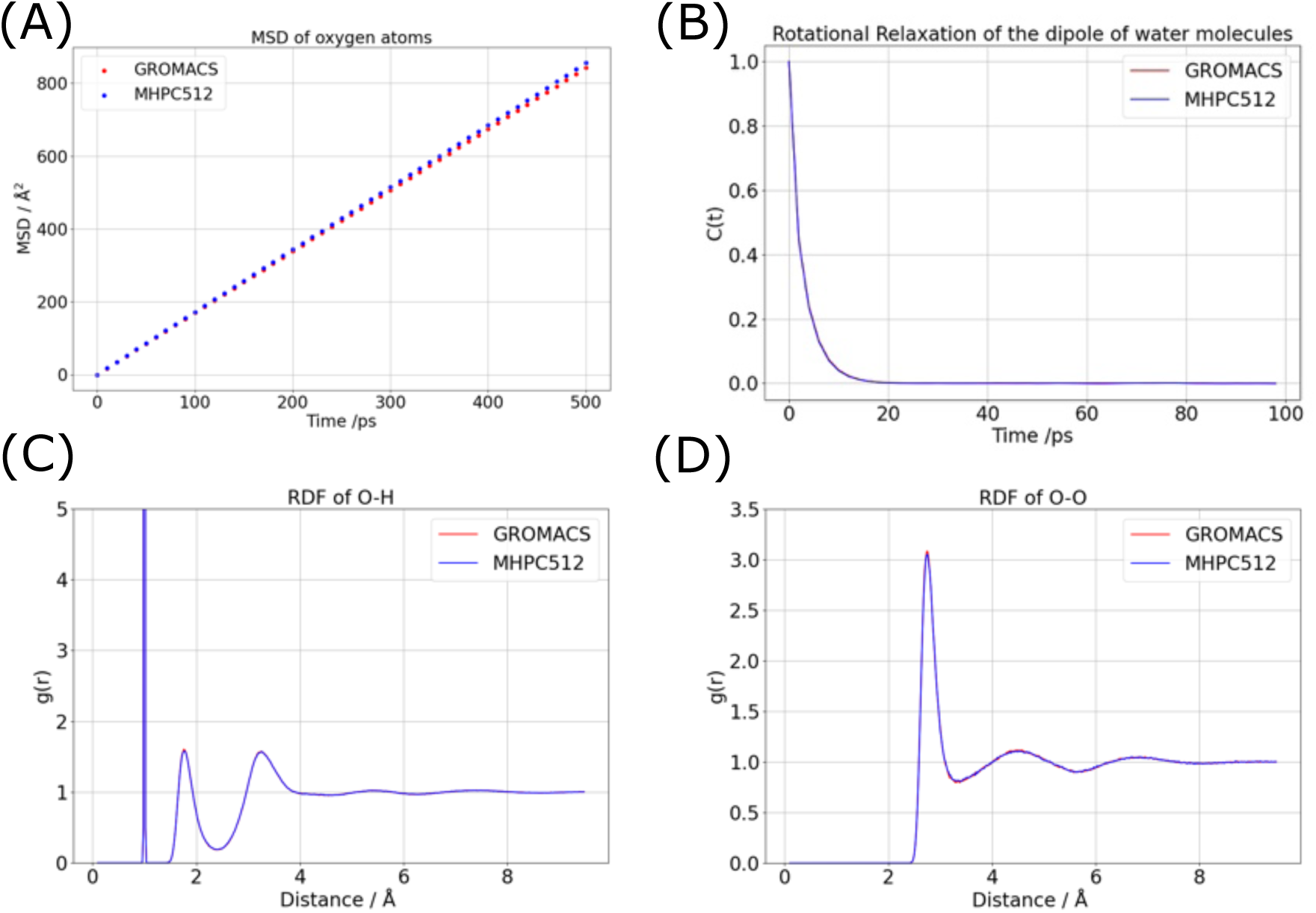
Structural characteristics of a SPC/E water box in a 100-ns simulation. (A) Mean square displacement (MSD) of the oxygen atoms. (B) Rotation relaxation of the dipole of water molecules. (C) Radial distribution function (RDF) of O – H. (D) RDF of O – O.

### 3.4 Dipeptides

The Ramachandran plots of dipeptides are commonly used to verify the reliability of force fields. However, here the Ramachandral plots of six dipeptides were used to verify whether the conformation distributions are consistent between MHPC512 and GROMACS. Molecular dynamics simulations were performed on amino acids in explicit water with capped terminals (acetyl and N-methylamide for the N-terminal and C-terminal, respectively). The simulation systems consisted of ACE-X-NME, where X represents Ala, Arg, Glu, Gly, Pro, and Trp.

The main chain dihedral angles of the central amino acids were obtained by analyzing the production molecular dynamics trajectories using MDAnalysis^68^. The resulting Ф and Ψ angles were mapped onto a 360 *×* 360 grid (with each coordinate having a grid width of 1*^°^*), and their distribution histogram was calculated. Shown in Fig. 5 are the natural logarithm of the histogram counts, reflecting the sampling of dihedral angles throughout the entire simulation time. The free energy landscapes shown in Fig. 5 are highly similar, indicating that MHPC512 can accurately sample the main chain dihedral angle distribution of proteins with a precision comparable to that of GROMACS. Quantitatively, by defining the secondary structure as helix if *-*180*^°^ <* Ф *<* 0*^°^* and *-*100*^°^ <* Ψ *<* 45*^°^*, PPII if *-*104*^°^ ≤ Ф ≤ -*46*^°^* and 116*^°^ ≤ Ψ ≤* 174*^°^*, and *ϑ* if *-*180*^°^ < Ф < -*45*^°^* and 45*^°^ < Ψ <* 225*^°^*, provided it is not classified as PPII. Using these definitions, the population in each dominant conformational region was counted. The results are presented in Table 3. It is evident that for all dipeptides, the populations of the three secondary structures differ by less than 1.2% between MHPC512 and GROMACS simulations, with the largest discrepancy observed in the PPII conformation of tryptophan.

**Figure 5:**
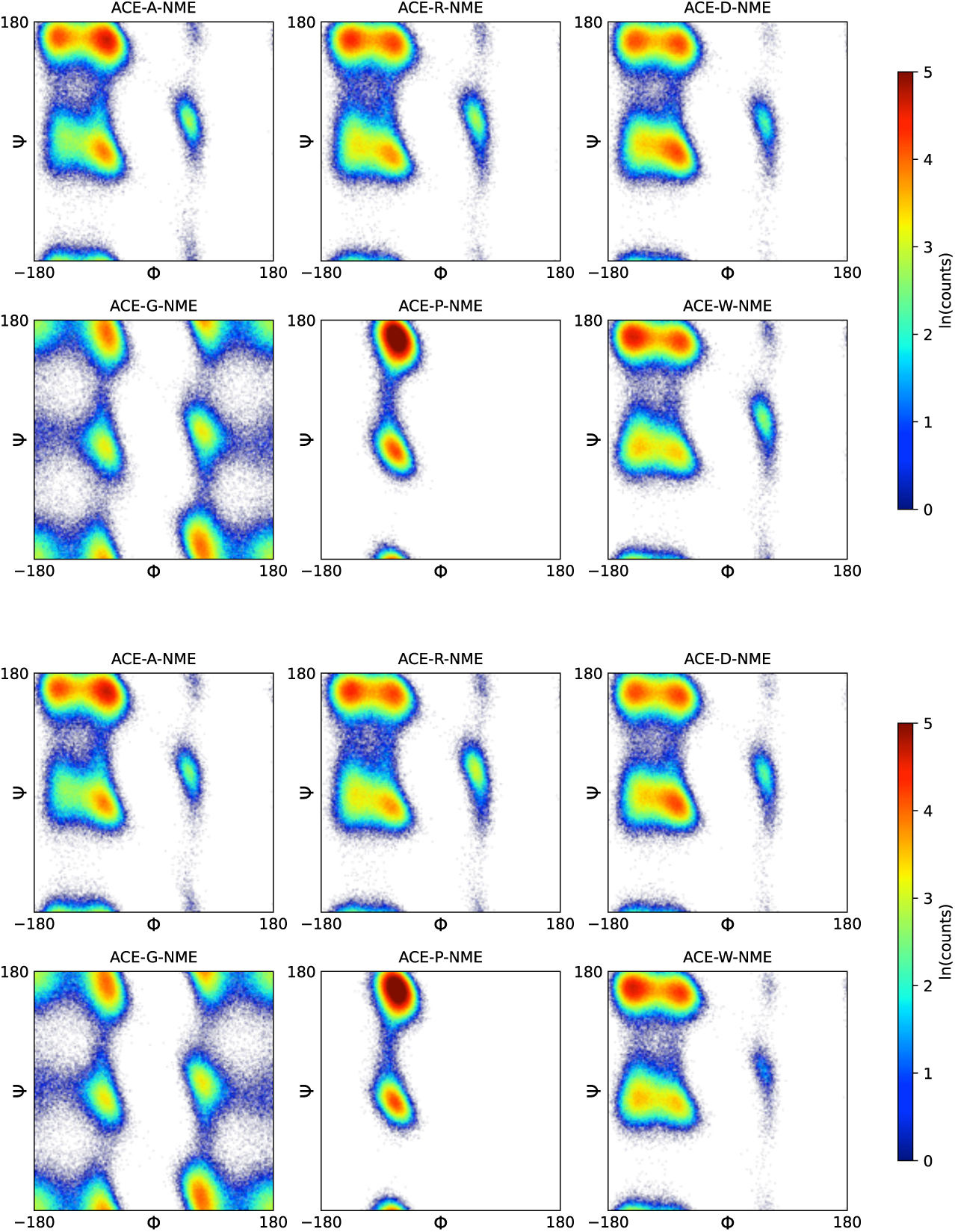
Ramachandran plots of dipeptides from the simulations of (top) MHPC512 and (bottom) GROMACS on Nvidia RTX 4090D.

**Table 3:** Secondary structure propensity for dipeptides (%)

|  |  | Ala | Arg | Glu | Gly | Pro | Trp |
| --- | --- | --- | --- | --- | --- | --- | --- |
| Helix | MHPC512 | 29.62 | 35.35 | 43.96 | 11.77 | 16.68 | 32.13 |
|  | GROMACS | 28.64 | 34.39 | 43.63 | 11.97 | 16.40 | 32.92 |
| $\beta$ | MHPC512 | 32.40 | 35.16 | 30.49 | 19.79 | 7.91 | 40.72 |
|  | GROMACS | 33.08 | 34.85 | 30.41 | 19.87 | 7.80 | 41.46 |
| PPII | MHPC512 | 33.65 | 24.49 | 22.86 | 16.88 | 73.30 | 23.18 |
|  | GROMACS | 34.59 | 24.72 | 23.00 | 16.91 | 73.77 | 24.29 |

### 3.5 Folding

Molecular dynamics simulations of protein folding processes serve as a means to validate the capabilities of a computing platform. In 2011, DESRES successfully simulated the folding of several short peptides using Anton.^69^ For the present work, three of these short peptides were selected, and their folding processes were simulated on MHPC512 using force fields consistent with those described in the original Science article.^69^ To track conformational changes during the simulations, two metrics were employed: the root mean square deviation (*C_α_*-RMSD) of all *C_α_* atoms relative to the experimental structure and the time evolution of secondary structure for each residue analyzed using the Define Secondary Structure of Protein (DSSP) method.^70^ The successful folding of these peptides, detailed below, demonstrates again that MHPC512 can reliably study long-time behaviors of peptides.

#### 3.5.1 Chignolin (CLN025)

Chignolin, a 10-residue peptide (sequence YYDPETGTWY, also known as CLN025), was chosen as the first test system. Figure 6 shows that Chignolin quickly found its folded state in both the MHPC512 and the GROMACS simulations with the *C_α_*-RMSD *<*2 Å from the NMR structure^71^. After that, both trajectories exhibited multiple folding–unfolding transitions. In the MHPC512 run, the peptide occupied the folded basin (*C_α_*-RMSD *<*2 Å from the NMR structure^71^) for 75.8% of the simulation time (Fig. 6). In the GROMACS run, the folded population was around 87.0%. The slight discrepancy may stem from the different treatments of long-range van der Waals interactions, yet overall MHPC512 reproduces the folding behavior of chignolin with comparable reliability to GROMACS.

**Figure 6:**
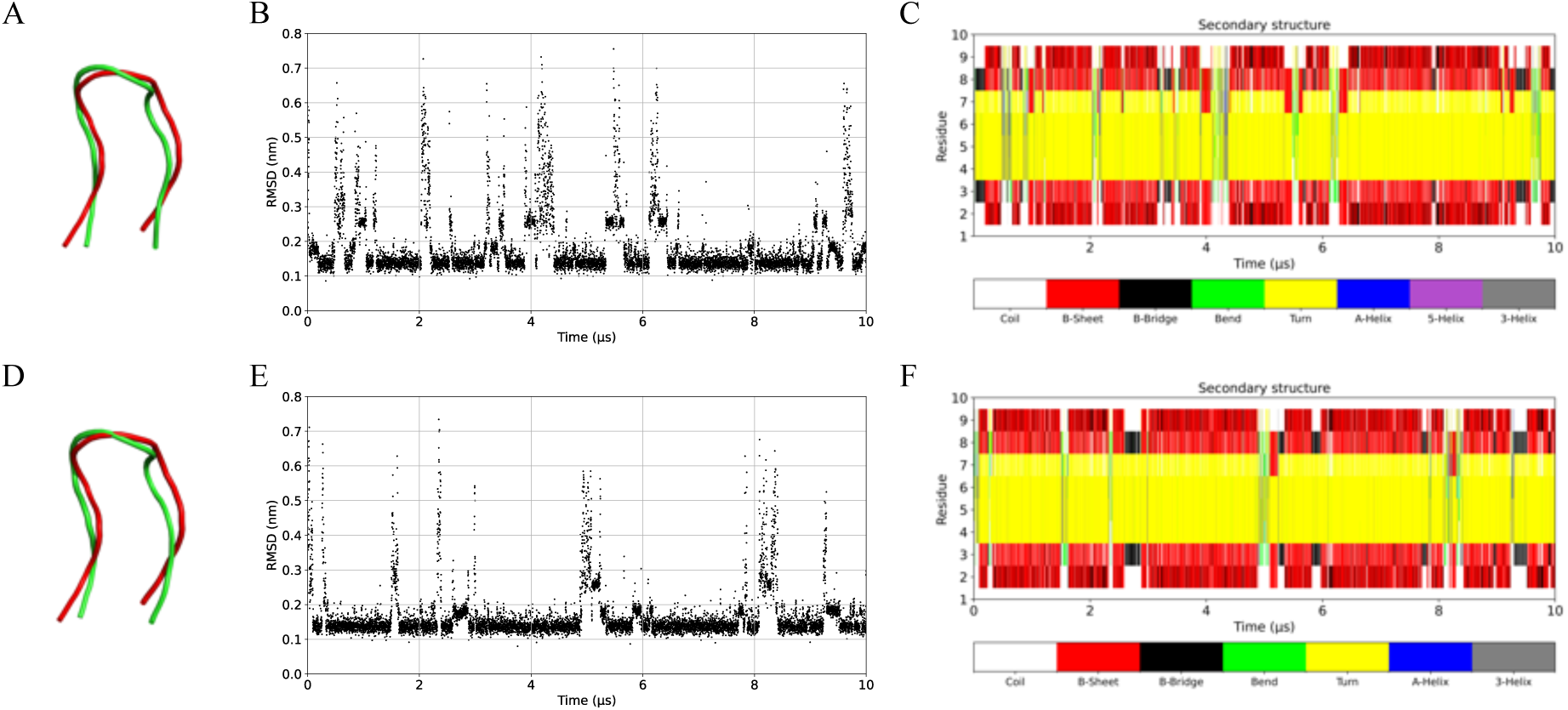
(A) Superposition of the folded structure (green, t =1.407 *µ*s with *C_α_* RMSD = 1.48 Å) and the experimental structure (red, PDB ID: 2RVD). The time evolution of (B) *C_α_*RMSD and (C) secondary structure during the MHPC simulation. (D) Superposition of the folded structure (t =8.461 *µ*s with *C_α_*RMSD = 1.83 Å) and the experimental structure (red). The time evolution of (E) *C_α_* RMSD and (F) secondary structure during the GROMACS simulation.

#### 3.5.2 Trp-cage

The second peptide for validation, Trp-cage, has been the subject of extensive folding studies with a variety of force fields.^72–80^ Trp-cage comprises 20 residues, of which residues 2–8 form an *ϖ*-helix and residues 12–14 form a short 3_10_ helix.^81^ The C*ϖ*-RMSD relative to the experimental structure and the residue-wise secondary-structure timeline are presented in Fig. 7. Starting from an extended chain, the peptide reached the folded state at around 9.5 *µ*s and remained folded for about 5 *µ*s before undergoing unfolding–refolding events. Although full convergence of the folded-state population was not achieved, the folding/unfolding behavior closely matches that reported by DESRES under the same force field, supporting the capability of MHPC512 to simulate Trp-cage folding accurately with CHARMM22*↑*.

**Figure 7:**
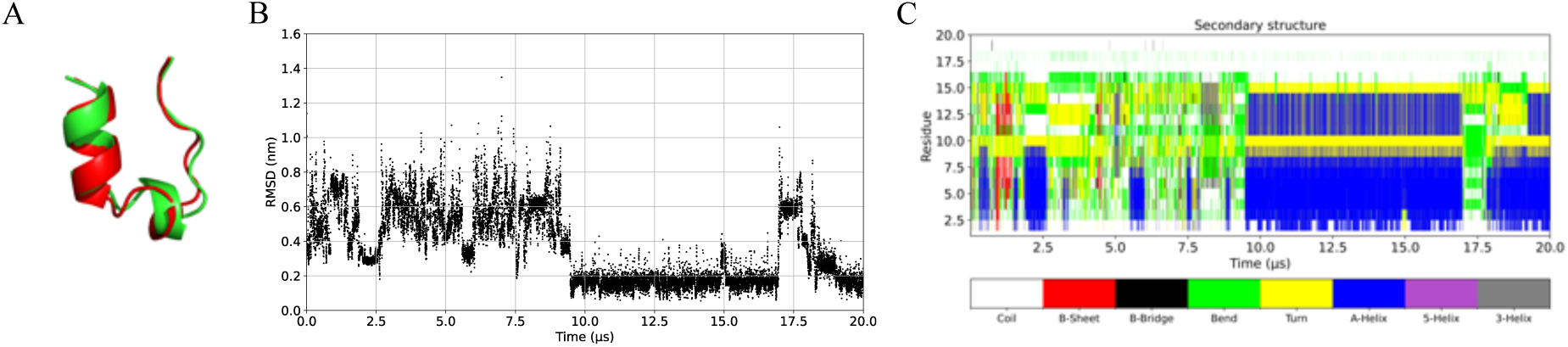
(A) Superposition of the folded structure (green, t =15.990 *µ*s with *C_α_* RMSD = 1.53 Å) and the experimental structure (red, PDB ID: 2JOF). The time evolution of (B) *C_α_*RMSD and (C) secondary structure during the MHPC simulation.

#### 3.5.3 Protein B

The native structure of Protein B consists of three helices that pack together. The representative folded structure, along with the time evolution of *C_α_*-RMSD relative to the experimental crystal structure and the corresponding secondary-structure profile, is presented in Fig. 8. The *C_α_*-RMSD indicates that it took over 10 *µ*s for this peptide to find its native conformation, although it briefly approached this structure around 7 *µ*s. The time evolution of secondary structures reveals that the helical regions fold rapidly, particularly the first two helices. However, the formation of the tertiary structure is slower. Throughout the trajectory, multiple folding and unfolding events were observed, with a stably folded ensemble persisting from 13 *µ*s to 22 *µ*s.

**Figure 8:**
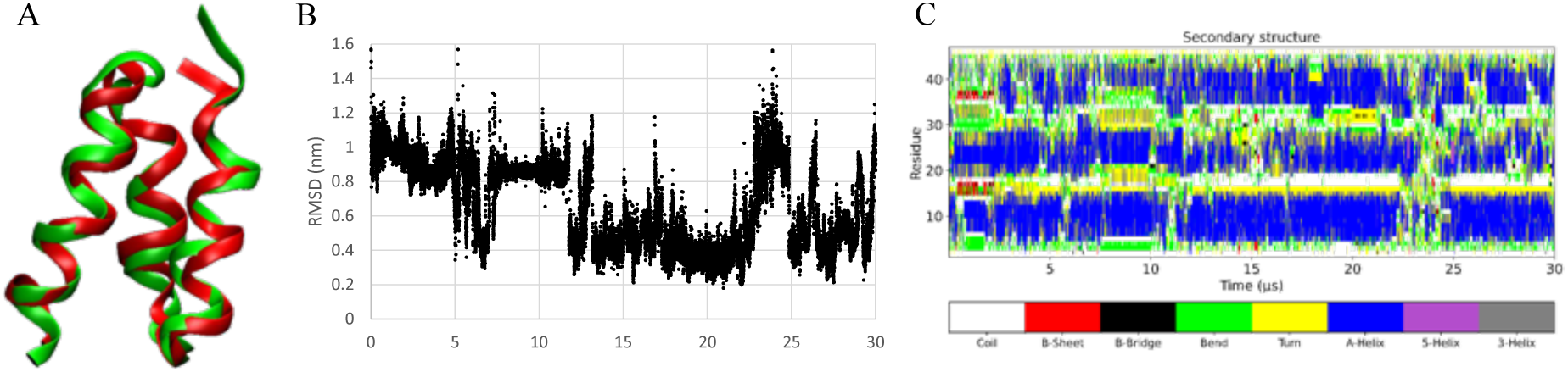
(A) Superposition of the folded structure (green, t = 20.948 *µ*s with *C_α_* RMSD = 1.99 Å) and the experimental structure of the K5I/K39V double mutant of the Protein B binding domain (red, PDB ID: 1PRB). The time evolution of (B) *C_α_* RMSD and (C) secondary structure during the MHPC512 simulation.

### 3.6 Membrane Systems

To assess the reliability of MHPC512 for membrane-protein simulations, the potassium channel KcsA from Streptomyces lividans (PDB ID 1BL8^82^) embedded in a POPC bilayer was simulated, and the results were compared with those obtained using GROMACS. Two force-fields—AMBER19SB/Lipid17 and CHARMM36m—were examined with TIP3P water model. Average box dimensions and backbone RMSD are used to measure the agreement between MHPC512 and Gromacs.

In the AMBER19SB + Lipid17 system, the average box lengths calculated by MHPC512 (10.06 nm *×* 9.98 nm *×* 8.46 nm) and by GROMACS (10.19 nm *×* 10.10 nm *×* 8.34 nm) differed by less than 0.15 nm in any dimension (see Fig. 9). Backbone RMSD profiles were also consistent, stabilizing at approximately 0.18 nm during the final 100 ns in both trajectories, indicating robust structural stability. For the CHARMM36m system, MHPC512 produced box dimensions of 10.24 nm *×* 10.24 nm *×* 8.70 nm, while GROMACS reported 10.39 nm *×* 10.39 nm *×* 8.54 nm (see Fig. 10). The backbone RMSD fluctuated around 0.10 nm in the last 100 ns for both simulations. No significant drift or anomalous behavior was observed at any stage. Taken together, these examinations demonstrate that MHPC512 reproduces membrane-protein simulations with accuracy comparable to GROMACS.

**Figure 9:**
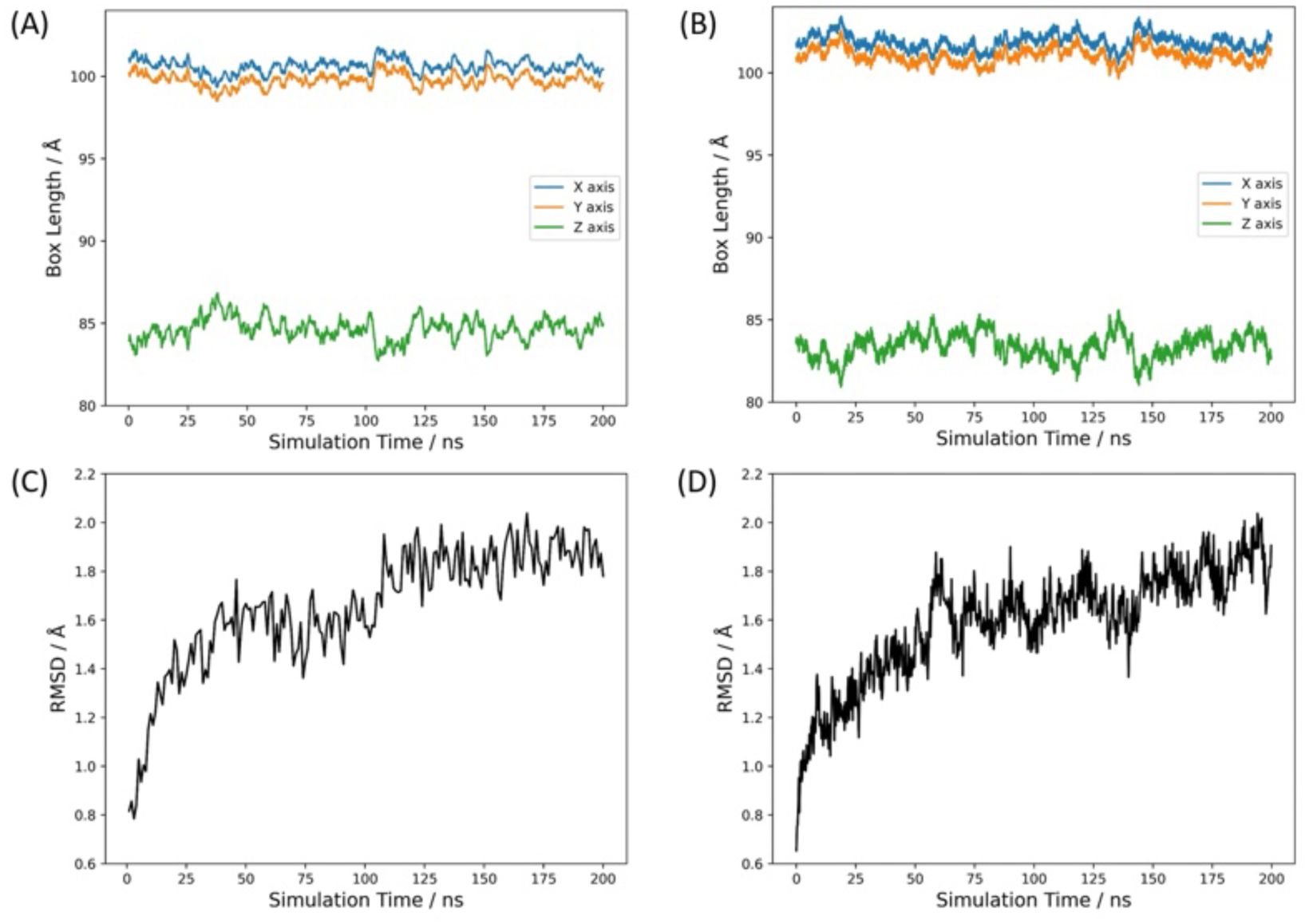
Evolution of the box dimensions during the (A) MHPC512 and (B) GROMACS simulations. Root mean square deviation of the protein backbone during the (C) MHPC512 and (D) GROMACS simulations. AMBER19SB and Lipid17 force fields are used for the protein and the lipid, respectively.

**Figure 10:**
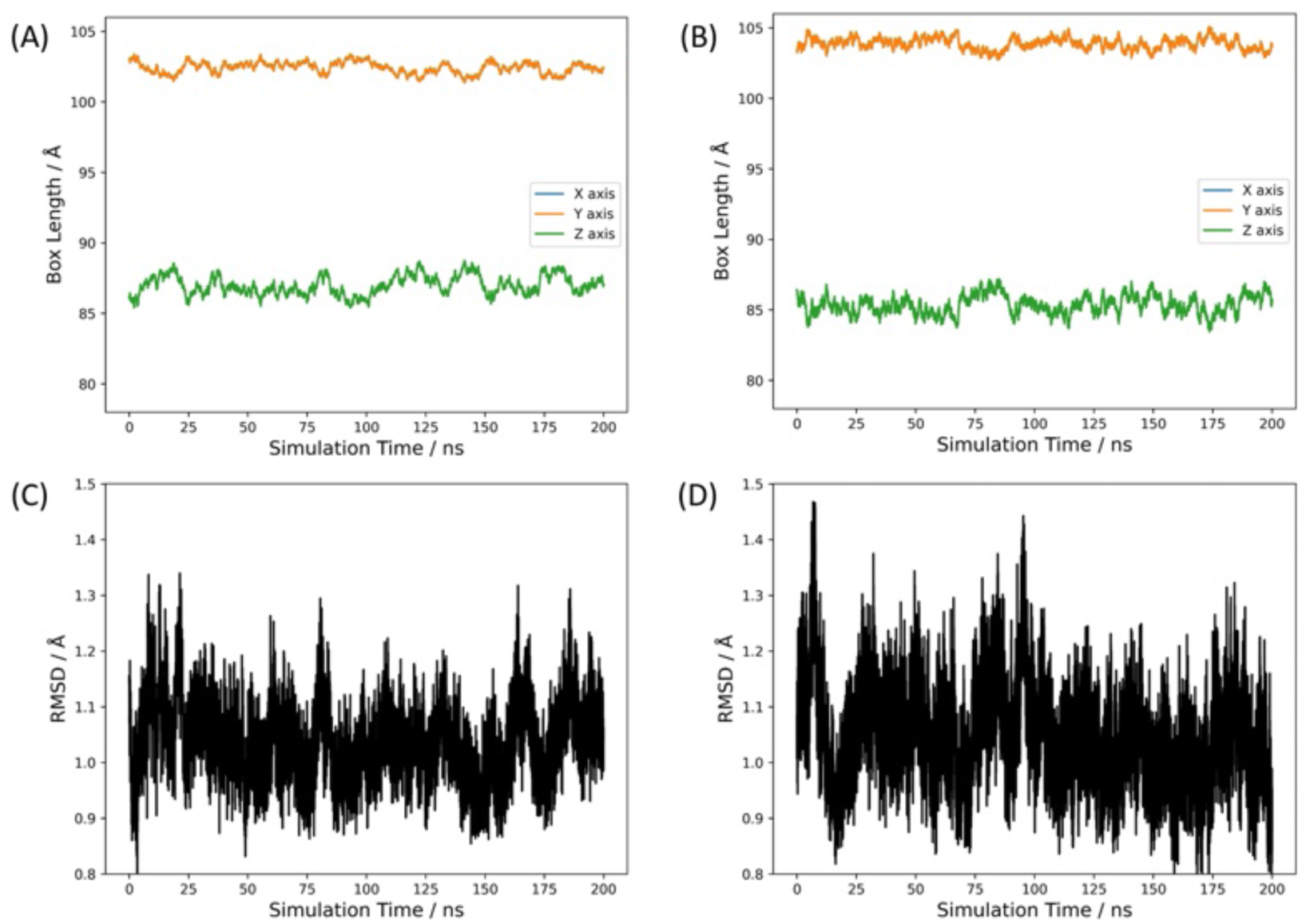
Evolution of the box dimensions during the (A) MHPC512 and (B) GROMACS simulations. Root mean square deviation of the protein backbone during the (C) MHPC512 and (D) GROMACS simulations. CHARMM36m force field is used for both the protein and the lipid.

## 4 Applications

### 4.1 Self Assembly of Lipid Molecules in Aqueous Solution

The initial structure is depicted in Fig. 11A, where lipid and water molecules were uniformly mixed within a box measuring 13.00 nm × 13.00 nm × 13.00 nm. As shown in Fig. 11B, isotropic compression during density relaxation caused structural distortions and reduced the box size to 9.74 nm × 9.74 nm × 9.74 nm. Although a complete bilayer had not yet formed, lipid molecules began to align with their nonpolar tails oriented inward and polar heads exposed to the surrounding water. Switching to semi-isotropic pressure coupling allowed the XY and Z dimensions to equilibrate independently. After 180 ns of semi-isotropic pressure coupling, the lipids continued to self-assemble, resulting in a box size of 10.11 nm × 10.11 nm × 8.94 nm. In this configuration, the Z dimension continued to shrink while the XY dimensions expanded, as illustrated in Fig. 11C. By the end of the 2-*µ*s simulation on MHPC512, the lipid molecules successfully self-assembled into a bilayer, and the box dimensions adjusted to 9.79 nm × 9.79 nm × 9.54 nm, as shown in Fig. 11D.

**Figure 11:**
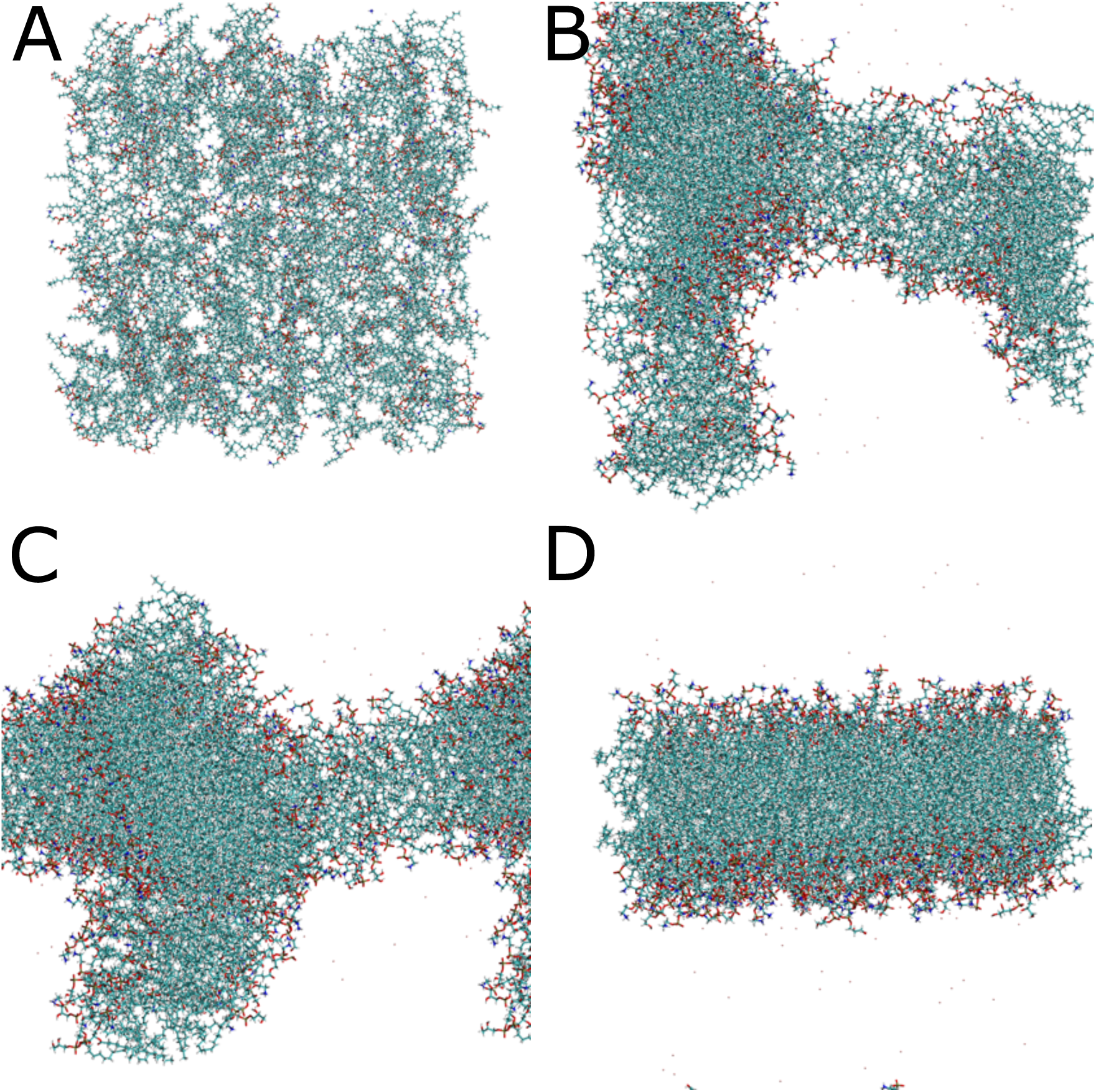
Evolution of the structures of the lipid aqueous solution during the simulation. (A) The initial structure. (B) The structure after the 20-ns isotropic NPT simulation. (C) The structure after 180-ns simulation on MHPC512. (D) The final structure after the 2-*µ*s simulation on MHPC512.

### 4.2 Conformation Transition between Alternative State and Canonical Active State for AT_1_R

Biased signaling, wherein distinct ligands engage G protein–coupled receptors (GPCRs) to selectively activate specific intracellular pathways, represents a promising strategy for developing safer and more efficacious therapeutics.^83^ The angiotensin II (AngII) type 1 receptor (AT_1_R) is a prototypical model for investigating the mechanism of biased signaling, where small modifications to AngII can result in either arrestin-biased or G protein–biased ligands and induce a higher or lower ratio of arrestin signaling to G protein signaling. ^84,85^ Following previous work^86^, a molecular dynamics simulation was initiated using the crystal structure of the active AT_1_R–AngII–NbAT110i1 complex (PDB ID: 6OS0) with the nanobody NbAT110i1 removed.

As illustrated in Fig. 12, the 5-*µ*s simulation reproduced both the alternative and canonical active conformations previously observed by Suomivuori et al.^86^ Both the alternative and canonical active conformations preserve the TM6 conformation observed in the nanobody-bound AT1R structure (PDB ID: 6OS0), which is characteristic of active-state GPCR structures. These two conformations show distinct intracellular arrangements, primarily involving TM7. In the alternative conformation, N1.50 loses its hydrogen bond with C7.47, which is re-established as a new hydrogen bond with N7.46 in the canonical active conformation. Additionally, the side chain of Y7.53 adopts a “downward” rotamer in the alternative conformation, oriented toward the intracellular side, whereas in the canonical active conformation, it switches to an “upward” rotamer pointing toward the extracellular side. The alternative conformation is proposed to preferentially couple with *ϑ*-arrestins, whereas the canonical active conformation is capable of engaging both Gq proteins and *ϑ*-arrestins. Transitions between these two conformational states have provided key mechanistic insights into the structural basis of biased signaling. In this 5-*µ*s simulation, this transition was also observed multiple times (Fig. 12). In total, these results demonstrate that MHPC512 is a practical and effective tool for studying GPCR dynamics and biased signaling mechanisms.

**Figure 12:**
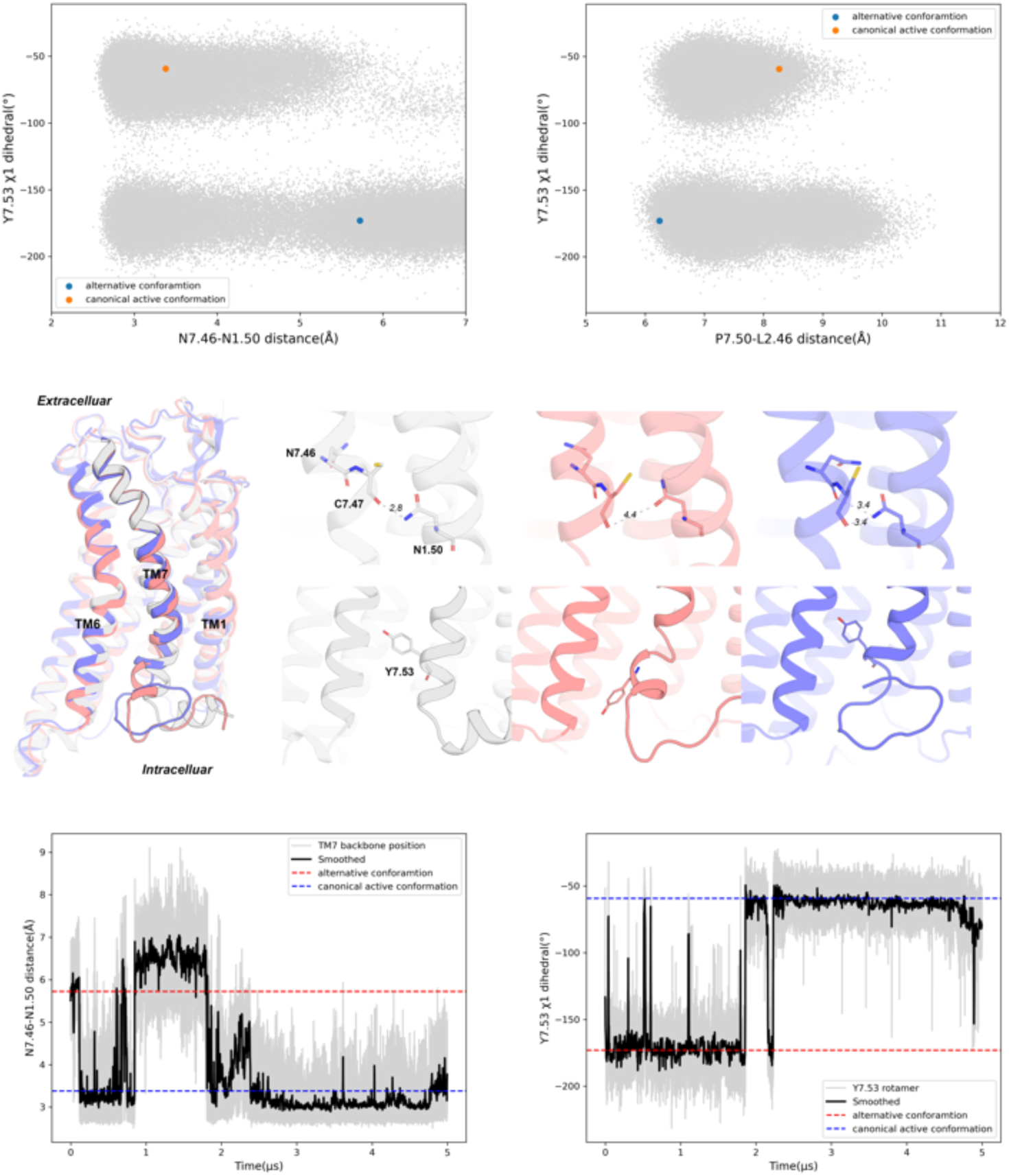
Top: Observed conformations of transmembrane helix 7 (TM7) in AT_1_R simulations projected to the 2D spaces characterized by the values of Y7.53 *ϱ*1 dihedral, N7.46-N1.50 O-ND2 distance and P7.50-L2.46 CA-CA distance. The orange and blue points represent reference structures corresponding to the alternative conformation and the canonical active conformation, respectively, provided in Ref. 86. Middle: Comparison between crystal structure and representative alternative conformation (1.817575 *µ*s) as well as canonical active conformation (4.360775 *µ*s). Bottom: Transition between alternative conformation and canonical active conformation. The grey line represents the unsmoothed traces, while the black line shows the traces smoothed using a moving average over a 2.5 ns window. The red and blue dash line represent reference structures corresponding to the alternative conformation and the canonical active conformation, respectively, provided in Ref. 86.

## 5 Future Perspective

As described above, MHPC512 currently supports standard molecular dynamics (MD) simulations using pairwise force fields and is publicly accessible to serve the community by accommodating many present-day simulation needs. Future updates will expand its capabilities in other applications such as material science (at both quantum mechanical and molecular mechanical levels of theory). While the current implementation enables simulations of systems with up to 1 million atoms over microsecond timescales, such durations may still be insufficient to capture many biologically relevant processes. To address this limitation, enhanced sampling techniques—such as temperature-space traversal (e.g., simulated tempering) and pulling along selected degrees of freedom—are planned for future integration. Incorporating more efficient constraint algorithms, including LINCS and SETTLE, is expected to improve computational performance. MHPC512 was originally designed for long-timescale production runs, and thus currently lacks several preparatory functions, such as energy minimization and large magnitude volume relaxation. Adding these features will boost global efficiency for a complete simulation (from the perspective of Amdahl’s argument), as preparing large systems (e.g., 1-million-atom models) remains time-consuming on general-purpose platforms. Finally, to better accommodate systems where electronic polarization plays a significant role, support for polarizable force fields is also under consideration for future development.

## References

(1) Karplus, M.; Petsko, G. A. Molecular Dynamics Simulations in Biology. Nature 1990, 347, 631–639.

(2) Allen, M. P.; Tildesley, D. J. Computer Simulation of Liquids; Oxford University Press, 2017.

(3) Andreoni, W.; Yip, S. Handbook of Materials Modeling; Springer, 2020.

(4) Pal, S.; Reddy, K. V. Molecular Dynamics for Materials Modeling: A Practical Approach Using LAMMPS Platform; CRC Press, 2024.

(5) Yip, S.; Short, M. P. Multiscale Materials Modelling at the Mesoscale. Nat. Mater. 2013, 12, 774–777.

(6) Gupta, C.; Sarkar, D.; Tieleman, D. P.; Singharoy, A. The Ugly, Bad, and Good Stories of Large-scale Biomolecular Simulations. Curr. Opin. Struct. Biol. 2022, 73, 102338.

(7) Ciccotti, G.; Dellago, C.; Ferrario, M.; Hernández, E.; Tuckerman, M. E. Molecular Simulations: Past, Present, and Future (A Topical Issue in EPJB). Eur. Phys. J. B 2022, 95, 3.

(8) Larsson, P.; Hess, B.; Lindahl, E. Algorithm Improvements for Molecular Dynamics Simulations. WIREs Comput. Mol. Sci. 2011, 1, 93–108.

(9) Hénin, J.; Lelièvre, T.; Shirts, M. R.; Valsson, O.; Delemotte, L. Enhanced Sampling Methods for Molecular Dynamics Simulations [Article v1.0]. Living J. Comp. Mol. Sci. 2022, 4, 1583.

(10) Keal, T. W.; Elena, A.-M.; Sokol, A. A.; Stoneham, K.; Probert, M. I. J.; Cucinotta, C. S.; Willock, D. J.; Logsdail, A. J.; Zen, A.; Hasnip, P. J.; Bush, I. J.; Watkins, M.; Alfè, D.; Skylaris, C.-K.; Curchod, B. F. E.; Cai, Q.; Woodley, S. M. Materials and Molecular Modeling at the Exascale. Comput. Sci. Eng. 2022, 24, 36–45.

(11) Stone, J. E.; Hardy, D. J.; Ufimtsev, I. S.; Schulten, K. GPU-accelerated Molecular Modeling Coming of Age. J. Mol. Graph. Model. 2010, 29, 116–125.

(12) Sugimoto, D.; Chikada, Y.; Makino, J.; Ito, T.; Ebisuzaki, T.; Umemura, M. A Special-purpose Computer for Gravitational Many-body Problems. Nature 1990, 345, 33–35.

(13) Ohmura, I.; Morimoto, G.; Ohno, Y.; Hasegawa, A.; Taiji, M. MDGRAPE-4: A Special-purpose Computer System for Molecular Dynamics Simulations. Philosophical Transactions of the Royal Society A: Mathematical, Physical and Engineering Sciences 2014, 372, 20130387.

(14) Shaw, D. E.; Deneroff, M. M.; Dror, R. O.; Kuskin, J. S.; Larson, R. H.; Salmon, J. K.; Young, C.; Batson, B.; Bowers, K. J.; Chao, J. C.; Eastwood, M. P.; Gagliardo, J.; Grossman, J. P.; Ho, C. R.; Ierardi, D. J.; Kolossváry, I.; Klepeis, J. L.; Layman, T.; McLeavey, C.; Moraes, M. A.; Mueller, R.; Priest, E. C.; Shan, Y.; Spengler, J.; Theobald, M.; Towles, B.; Wang, S. C. Anton, a Special-purpose Machine for Molecular Dynamics Simulation. Commun. ACM 2008, 51, 91–97.

(15) Shaw, D. E.; Grossman, J. P.; Bank, J. A.; Batson, B.; Butts, J. A.; Chao, J. C.; Deneroff, M. M.; Dror, R. O.; Even, A.; Fenton, C. H.; Forte, A.; Gagliardo, J.; Gill, G.; Greskamp, B.; Ho, C. R.; Ierardi, D. J.; Iserovich, L.; Kuskin, J. S.; Larson, R. H.; Layman, T.; Lee, L.-S.; Lerer, A. K.; Li, C.; Killebrew, D.; Mackenzie, K. M.; Mok, S. Y.-H.; Moraes, M. A.; Mueller, R.; Nociolo, L. J.; Peticolas, J. L.; Quan, T.; Ramot, D.; Salmon, J. K.; Scarpazza, D. P.; Ben Schafer, U.; Siddique, N.; Snyder, C. W.; Spengler, J.; Tang, P. T. P.; Theobald, M.; Toma, H.; Towles, B.; Vitale, B.; Wang, S. C.; Young, C. Anton 2: Raising the Bar for Performance and Programmability in a Special-purpose Molecular Dynamics Supercomputer. Proceedings of the International Conference for High Performance Computing, Networking, Storage and Analysis. 2014; p 41–53.

16. Shaw, D. E.; Adams, P. J.; Azaria, A.; Bank, J. A.; Batson, B.; Bell, A.; Bergdorf, M.; Bhatt, J.; Butts, J. A.; Correia, T.; Dirks, R. M.; Dror, R. O.; Eastwood, M. P.; Edwards, B.; Even, A.; Feldmann, P.; Fenn, M.; Fenton, C. H.; Forte, A.; Gagliardo, J.; Gill, G.; Gorlatova, M.; Greskamp, B.; Grossman, J.; Gullingsrud, J.; Harper, A.; Hasenplaugh, W.; Heily, M.; Heshmat, B. C.; Hunt, J.; Ierardi, D. J.; Iserovich, L.; Jackson, B. L.; Johnson, N. P.; Kirk, M. M.; Klepeis, J. L.; Kuskin, J. S.; Mackenzie, K. M.; Mader, R. J.; McGowen, R.; McLaughlin, A.; Moraes, M. A.; Nasr, M. H.; Nociolo, L. J.; O’Donnell, L.; Parker, A.; Peticolas, J. L.; Pocina, G.; Predescu, C.; Quan, T.; Salmon, J. K.; Schwink, C.; Shim, K. S.; Siddique, N.; Spengler, J.; Szalay, T.; Tabladillo, R.; Tartler, R.; Taube, A. G.; Theobald, M.; Towles, B.; Vick, W.; Wang, S. C.; Wazlowski, M.; Weingarten, M. J.; Williams, J. M.; Yuh, K. A. Anton 3: Twenty Microseconds of Molecular Dynamics Simulation before Lunch. Proceedings of the International Conference for High Performance Computing, Networking, Storage and Analysis. New York, NY, USA, 2021.

(17) Wang, D.; Du, X.; Yin, L.; Lin, C.; Ma, H.; Ren, W.; Wang, H.; Wang, X.; Xie, S.; Wang, L.; Liu, Z.; Wang, T.; Pu, Z.; Ding, G.; Zhu, M.; Yang, L.; Guo, R.; Zhang, Z.; Lin, X.; Hao, J.; Yang, Y.; Sun, W.; Zhou, F.; Xiao, N.; Cui, Q.; Wang, X. MaPU: A Novel Mathematical Computing Architecture. 2016 IEEE International Symposium on High Performance Computer Architecture (HPCA). 2016; pp 457–468.

(18) Abraham, M. J.; Murtola, T.; Schulz, R.; Páll, S.; Smith, J. C.; Hess, B.; Lindahl, E. GROMACS: High Performance Molecular Simulations through Multi-level Parallelism from Laptops to Supercomputers. SoftwareX 2015, 1-2, 19–25.

(19) Case, D. A.; Cheatham III, T. E.; Darden, T.; Gohlke, H.; Luo, R.; Merz Jr., K. M.; Onufriev, A.; Simmerling, C.; Wang, B.; Woods, R. J. The Amber Biomolecular Simulation Programs. J. Comput. Chem. 2005, 26, 1668–1688.

(20) Case, D. A.; Cerutti, D. S.; Cruzeiro, V. W. D.; Darden, T. A.; Duke, R. E.; Ghazimirsaeed, M.; Giambaşu, G. M.; Giese, T. J.; Götz, A. W.; Harris, J. A.; Kasavajhala, K.; Lee, T.-S.; Li, Z.; Lin, C.; Liu, J.; Miao, Y.; Salomon-Ferrrer, R.; Shen, J.; Snyder, R.; Swails, J.; Walker, R. C.; Wang, J.; Wu, X.; Zeng, J.; Cheatham III, T. E.; Roe, D. R.; Roitberg, A.; Simmerling, C.; York, D. M.; Nagan, M. C.; Merz, K. M. J. Recent Developments in Amber Biomolecular Simulations. J. Chem. Info. Model. 2025, 65, 7835–7843.

(21) Hwang, W.; Austin, S. L.; Blondel, A.; Boittier, E. D.; Boresch, S.; Buck, M.; Buckner, J.; Caflisch, A.; Chang, H.-T.; Cheng, X.; Choi, Y. K.; Chu, J.-W.; Crowley, M. F.; Cui, Q.; Damjanovic, A.; Deng, Y.; Devereux, M.; Ding, X.; Feig, M. F.; Gao, J.; Glowacki, D. R.; Gonzales, J. E. I.; Hamaneh, M. B.; Harder, E. D.; Hayes, R. L.; Huang, J.; Huang, Y.; Hudson, P. S.; Im, W.; Islam, S. M.; Jiang, W.; Jones, M. R.; Käser, S.; Kearns, F. L.; Kern, N. R.; Klauda, J. B.; Lazaridis, T.; Lee, J.; Lemkul, J. A.; Liu, X.; Luo, Y.; MacKerell, A. D. J.; Major, D. T.; Meuwly, M.; Nam, K.; Nilsson, L.; Ovchinnikov, V.; Paci, E.; Park, S.; Pastor, R. W.; Pittman, A. R.; Post, C. B.; Prasad, S.; Pu, J.; Qi, Y.; Rathinavelan, T.; Roe, D. R.; Roux, B.; Rowley, C. N.; Shen, J.; Simmonett, A. C.; Sodt, A. J.; Töpfer, K.; Upadhyay, M.; van der Vaart, A.; Vazquez-Salazar, L. I.; Venable, R. M.; Warrensford, L. C.; Woodcock, H. L.; Wu, Y.; Brooks, C. L. I.; Brooks, B. R.; Karplus, M. CHARMM at 45: Enhancements in Accessibility, Functionality, and Speed. J. Phys. Chem. B 2024, 128, 9976–10042.

(22) Kollman, P.; Dixon, R.; Cornell, W.; Fox, T.; Chipot, C.; Pohorille, A. In Computer Simulation of Biomolecular Systems: Theoretical and Experimental Applications; van Gunsteren, W. F., Weiner, P. K., Wilkinson, A. J., Eds.; Springer Netherlands: Dordrecht, 1997; pp 83–96.

(23) Jorgensen, W. L.; Maxwell, D. S.; Tirado-Rives, J. Development and Testing of the OPLS All-Atom Force Field on Conformational Energetics and Properties of Organic Liquids. J. Am. Chem. Soc. 1996, 118, 11225–11236.

(24) Bowers, K. J.; Dror, R. O.; Shaw, D. E. The Midpoint Method for Parallelization of Particle Simulations. J. Chem. Phys. 2006, 124, 184109.

(25) Bowers, K. J.; Dror, R. O.; Shaw, D. E. Zonal Methods for the Parallel Execution of Range-limited N-body Simulations. J. Comput. Phys. 2007, 221, 303–329.

(26) Shan, Y.; Klepeis, J. L.; Eastwood, M. P.; Dror, R. O.; Shaw, D. E. Gaussian Split Ewald: A Fast Ewald Mesh Method for Molecular Simulation. J. Chem. Phys. 2005, 122, 054101.

(27) Tuckerman, M.; Berne, B. J.; Martyna, G. J. Reversible Multiple Time Scale Molecular Dynamics. J. Chem. Phys. 1992, 97, 1990–2001.

(28) Trotter, H. F. On the Product of Semi-Groups of Operators. Proc. Am. Math. Soc. 1959, 10, 545–551.

(29) Lambrakos, S.; Boris, J.; Oran, E.; Chandrasekhar, I.; Nagumo, M. A Modified SHAKE Algorithm for Maintaining Rigid Bonds in Molecular Dynamics Simulations of Large Molecules. J. Comput. Phys. 1989, 85, 473–486.

(30) Andersen, H. C. Rattle: A “Velocity” Version of the SHAKE Algorithm for Molecular Dynamics Calculations. J. Comput. Phys. 1983, 52, 24–34.

(31) Berendsen, H. J. C.; Postma, J. P. M.; van Gunsteren, W. F.; DiNola, A.; Haak, J. R. Molecular Dynamics with Coupling to an External Bath. J. Chem. Phys. 1984, 81, 3684–3690.

(32) Martyna, G. J.; Klein, M. L.; Tuckerman, M. Nosé–Hoover Chains: The Canonical Ensemble via Continuous Dynamics. J. Chem. Phys. 1992, 97, 2635–2643.

(33) Parrinello, M.; Rahman, A. Polymorphic Transitions in Single Crystals: A New Molecular Dynamics Method. J. Appl. Phys. 1981, 52, 7182–7190.

(34) Andersen, H. C. Molecular Dynamics Simulations at Constant Pressure and/or Temperature. J. Chem. Phys. 1980, 72, 2384–2393.

(35) Zhang, Y.; Feller, S. E.; Brooks, B. R.; Pastor, R. W. Computer Simulation of Liquid/Liquid Interfaces. I. Theory and Application to Octane/Water. J. Chem. Phys. 1995, 103, 10252–10266.

(36) Case, D. A.; Aktulga, H. M.; Belfon, K.; Cerutti, D. S.; Cisneros, G. A.; Cruzeiro, V. W. D.; Forouzesh, N.; Giese, T. J.; Götz, A. W.; Gohlke, H.; Izadi, S.; Kasavajhala, K.; Kaymak, M. C.; King, E.; Kurtzman, T.; Lee, T.-S.; Li, P.; Liu, J.; Luchko, T.; Luo, R.; Manathunga, M.; Machado, M. R.; Nguyen, H. M.; O’Hearn, K. A.; Onufriev, A. V.; Pan, F.; Pantano, S.; Qi, R.; Rahnamoun, A.; Risheh, A.; Schott-Verdugo, S.; Shajan, A.; Swails, J.; Wang, J.; Wei, H.; Wu, X.; Wu, Y.; Zhang, S.; Zhao, S.; Zhu, Q.; Cheatham, T. E. I.; Roe, D. R.; Roitberg, A.; Simmerling, C.; York, D. M.; Nagan, M. C.; Merz, K. M. J. AmberTools. J. Chem. Info. Model. 2023, 63, 6183–6191.

(37) Robertson, M. J.; Tirado-Rives, J.; Jorgensen, W. L. Improved Peptide and Protein Torsional Energetics with the OPLS-AA Force Field. J. Chem. Theory Comput. 2015, 11, 3499–3509.

(38) Robertson, M. J.; Qian, Y.; Robinson, M. C.; Tirado-Rives, J.; Jorgensen, W. L. Development and Testing of the OPLS-AA/M Force Field for RNA. J. Chem. Theory Comput. 2019, 15, 2734–2742.

(39) Piana, S.; Lindorff-Larsen, K.; Shaw, D. How Robust Are Protein Folding Simulations with Respect to Force Field Parameterization? Biophys. J. 2011, 100, L47–L49.

(40) Mackerell Jr., A. D.; Feig, M.; Brooks III, C. L. Extending the Treatment of Backbone Energetics in Protein Force Fields: Limitations of Gas-phase Quantum Mechanics in Reproducing Protein Conformational Distributions in Molecular Dynamics Simulations. J. Comput. Chem. 2004, 25, 1400–1415.

(41) Huang, J.; MacKerell Jr, A. D. CHARMM36 All-atom Additive Protein Force Field: Validation Based on Comparison to NMR Data. J. Comput. Chem. 2013, 34, 2135–2145.

(42) Huang, J.; Rauscher, S.; Nawrocki, G.; Ran, T.; Feig, M.; De Groot, B. L.; Grubmüller, H.; MacKerell Jr, A. D. CHARMM36m: An Improved Force Field for Folded and Intrinsically Disordered Proteins. Nat. Methods 2017, 14, 71–73.

(43) Lindorff-Larsen, K.; Piana, S.; Palmo, K.; Maragakis, P.; Klepeis, J. L.; Dror, R. O.; Shaw, D. E. Improved Side-chain Torsion Potentials for the Amber ff99SB Protein Force Field. *Proteins: Struct., Funct.*, Bioinf. 2010, 78, 1950–1958.

(44) Maier, J. A.; Martinez, C.; Kasavajhala, K.; Wickstrom, L.; Hauser, K. E.; Simmerling, C. ff14SB: Improving the Accuracy of Protein Side Chain and Backbone Parameters from ff99SB. J. Chem. Theory Comput. 2015, 11, 3696–3713.

(45) Tian, C.; Kasavajhala, K.; Belfon, K. A. A.; Raguette, L.; Huang, H.; Migues, A. N.; Bickel, J.; Wang, Y.; Pincay, J.; Wu, Q.; Simmerling, C. ff19SB: Amino-Acid-Specific Protein Backbone Parameters Trained against Quantum Mechanics Energy Surfaces in Solution. J. Chem. Theory Comput. 2020, 16, 528–552.

(46) Wang, J.; Wolf, R. M.; Caldwell, J. W.; Kollman, P. A.; Case, D. A. Development and Testing of a General Amber Force Field. J. Comput. Chem. 2004, 25, 1157–1174.

(47) Dickson, C. J.; Walker, R. C.; Gould, I. R. Lipid21: Complex Lipid Membrane Simulations with AMBER. J. Chem. Theory Comput. 2022, 18, 1726–1736.

(48) Ivani, I.; Dans, P. D.; Noy, A.; Pérez, A.; Faustino, I.; Hospital, A.; Walther, J.; Andrio, P.; Goñi, R.; Balaceanu, A.; others Parmbsc1: A Refined Force Field for DNA Simulations. Nat. Methods 2016, 13, 55–58.

(49) Zgarbová, M.; Šponer, J.; Otyepka, M.; Cheatham, T. E. I.; Galindo-Murillo, R.; Jurečka, P. Refinement of the Sugar–Phosphate Backbone Torsion Beta for AMBER Force Fields Improves the Description of Z- and B-DNA. J. Chem. Theory Comput. 2015, 11, 5723–5736.

(50) Zgarbová, M.; Šponer, J.; Jurečka, P. Z-DNA as a Touchstone for Additive Empirical Force Fields and a Refinement of the Alpha/Gamma DNA Torsions for AMBER. J. Chem. Theory Comput. 2021, 17, 6292–6301.

(51) Zgarbová, M.; Šponer, J.; Jurečka, P. Refinement of the Sugar Puckering Torsion Potential in the AMBER DNA Force Field. J. Chem. Theory Comput. 2025, 21, 833–846.

(52) Liebl, K.; Zacharias, M. Tumuc1: A New Accurate DNA Force Field Consistent with High-Level Quantum Chemistry. J. Chem. Theory Comput. 2021, 17, 7096–7105.

(53) Zgarbová, M.; Otyepka, M.; Šponer, J.; Mládek, A.; Banáš, P.; Cheatham III, T. E.; Jurev̌ka, P. Refinement of the Cornell et al. Nucleic Acids Force Field Based on Reference Quantum Chemical Calculations of Glycosidic Torsion Profiles. J. Chem. Theory Comput. 2011, 7, 2886–2902.

(54) Bergonzo, C.; Cheatham III, T. E. Improved Force Field Parameters Lead to a Better Description of RNA Structure. J. Chem. Theory Comput. 2015, 11, 3969–3972.

(55) Tan, D.; Piana, S.; Dirks, R. M.; Shaw, D. E. RNA Force Field with Accuracy Comparable to State-of-the-art Protein Force Fields. Proc. Natl. Acad. Sci. U.S.A. 2018, 115, E1346–E1355.

(56) Kirschner, K. N.; Yongye, A. B.; Tschampel, S. M.; González-Outeiriño, J.; Daniels, C. R.; Foley, B. L.; Woods, R. J. GLYCAM06: A Generalizable Biomolecular Force Field. Carbohydrates. J. Comput. Chem. 2008, 29, 622–655.

(57) Homeyer, N.; Horn, A. H.; Lanig, H.; Sticht, H. AMBER Force-field Parameters for Phosphorylated Amino Acids in Different Protonation States: Phosphoserine, Phospho-threonine, Phosphotyrosine, and Phosphohistidine. J. Mol. Model. 2006, 12, 281–289.

(58) Raguette, L. E.; Cuomo, A. E.; Belfon, K. A. A.; Tian, C.; Hazoglou, V.; Witek, G.; Telehany, S. M.; Wu, Q.; Simmerling, C. phosaa14SB and phosaa19SB: Updated Amber Force Field Parameters for Phosphorylated Amino Acids. J. Chem. Theory Comput. 2024, 20, 7199–7209.

(59) Jorgensen, W. L.; Chandrasekhar, J.; Madura, J. D.; Impey, R. W.; Klein, M. L. Comparison of Simple Potential Functions for Simulating Liquid Water. J. Chem. Phys. 1983, 79, 926–935.

(60) MacKerell Jr., A. D.; Bashford, D.; Bellott, M.; Dunbrack, R. L. J.; Evanseck, J. D.; Field, M. J.; Fischer, S.; Gao, J.; Guo, H.; Ha, S.; Joseph-McCarthy, D.; Kuchnir, L.; Kuczera, K.; Lau, F. T. K.; Mattos, C.; Michnick, S.; Ngo, T.; Nguyen, D. T.; Prodhom, B.; Reiher, W. E.; Roux, B.; Schlenkrich, M.; Smith, J. C.; Stote, R.; Straub, J.; Watanabe, M.; Wiórkiewicz-Kuczera, J.; Yin, D.; Karplus, M. All-Atom Empirical Potential for Molecular Modeling and Dynamics Studies of Proteins. J. Phys. Chem. B 1998, 102, 3586–3616.

(61) Berendsen, H. J. C.; Postma, J. P. M.; van Gunsteren, W. F.; Hermans, J. In Intermolecular Forces: Proceedings of the Fourteenth Jerusalem Symposium on Quantum Chemistry and Biochemistry Held in Jerusalem, Israel, April 13–16, 1981; Pullman, B., Ed.; Springer Netherlands: Dordrecht, 1981; pp 331–342.

(62) Berweger, C. D.; van Gunsteren, W. F.; Müller-Plathe, F. Force Field Parametrization by Weak Coupling. Re-engineering SPC Water. Chem. Phys. Lett. 1995, 232, 429–436.

(63) Berendsen, H. J. C.; Grigera, J. R.; Straatsma, T. P. The Missing Term in Effective Pair Potentials. J. Phys. Chem. 1987, 91, 6269–6271.

(64) Izadi, S.; Anandakrishnan, R.; Onufriev, A. V. Building Water Models: A Different Approach. J. Phys. Chem. Lett. 2014, 5, 3863–3871.

(65) Wu, Y.; Tepper, H. L.; Voth, G. A. Flexible Simple Point-charge Water Model with Improved Liquid-State Properties. J. Chem. Phys. 2006, 124, 024503.

(66) Abraham, B. G.; Haikarainen, T.; Vuorio, J.; Girych, M.; Virtanen, A. T.; Kurttila, A.; Karathanasis, C.; Heilemann, M.; Shamrma, V.; Vattulainen, I.; Silvennoinen, O. Molecular Basis of JAK2 Activation in Erythropoietin Receptor and Pathogenic JAK2 Signaling. Sci. Adv. 2024, 10, eadl2097.

(67) Miyamoto, S.; Kollman, P. A. Settle: An Analytical Version of the SHAKE and RATTLE Algorithm for Rigid Water Models. J. Comput. Chem. 1992, 13, 952–962.

(68) Michaud-Agrawal, N.; Denning, E. J.; Woolf, T. B.; Beckstein, O. MDAnalysis: A Toolkit for the Analysis of Molecular Dynamics Simulations. J. Comput. Chem. 2011, 32, 2319–2327.

(69) Lindorff-Larsen, K.; Piana, S.; Dror, R. O.; Shaw, D. E. How Fast-Folding Proteins Fold. Science 2011, 334, 517–520.

(70) Kabsch, W.; Sander, C. Dictionary of Protein Secondary Structure: Pattern Recognition of Hydrogen-bonded and Geometrical Features. Biopolymers 1983, 22, 2577–2637.

(71) Honda, S.; Akiba, T.; Kato, Y. S.; Sawada, Y.; Sekijima, M.; Ishimura, M.; Ooishi, A.; Watanabe, H.; Odahara, T.; Harata, K. Crystal Structure of a Ten-Amino Acid Protein. J. Am. Chem. Soc. 2008, 130, 15327–15331.

(72) Zhou, R. Trp-cage: Folding Free Energy Landscape in Explicit Water. Proc. Natl. Acad. Sci. U.S.A. 2003, 100, 13280–13285.

(73) Pitera, J. W.; Swope, W. Understanding Folding and Design: Replica-exchange Simulations of “Trp-cage” Miniproteins. Proc. Natl. Acad. Sci. U.S.A. 2003, 100, 7587–7592.

(74) Marinelli, F.; Pietrucci, F.; Laio, A.; Piana, S. A Kinetic Model of Trp-Cage Folding from Multiple Biased Molecular Dynamics Simulations. PLOS Comput. Biol. 2009, 5, 1–18.

(75) Day, R.; Paschek, D.; Garcia, A. E. Microsecond Simulations of the Folding/Unfolding Thermodynamics of the Trp-cage Miniprotein. *Proteins: Struct., Funct.*, Bioinf. 2010, 78, 1889–1899.

(76) Shao, Q.; Shi, J.; Zhu, W. Enhanced Sampling Molecular Dynamics Simulation Captures Experimentally Suggested Intermediate and Unfolded States in the Folding Pathway of Trp-cage Miniprotein. J. Chem. Phys. 2012, 137, 125103.

(77) Marinelli, F. Following Easy Slope Paths on a Free Energy Landscape: The Case Study of the Trp-Cage Folding Mechanism. Biophys. J. 2013, 105, 1236–1247.

(78) Meuzelaar, H.; Marino, K. A.; Huerta-Viga, A.; Panman, M. R.; Smeenk, L. E. J.; Kettelarij, A. J.; van Maarseveen, J. H.; Timmerman, P.; Bolhuis, P. G.; Woutersen, S. Folding Dynamics of the Trp-Cage Miniprotein: Evidence for a Native-Like Intermediate from Combined Time-Resolved Vibrational Spectroscopy and Molecular Dynamics Simulations. J. Phys. Chem. B 2013, 117, 11490–11501.

(79) English, C. A.; García, A. E. Folding and Unfolding Thermodynamics of the TC10b Trp-cage Miniprotein. Phys. Chem. Chem. Phys. 2014, 16, 2748–2757.

(80) Kim, S. B.; Dsilva, C. J.; Kevrekidis, I. G.; Debenedetti, P. G. Systematic Characterization of Protein Folding Pathways Using Diffusion Maps: Application to Trp-cage Miniprotein. J. Chem. Phys. 2015, 142, 085101.

(81) Barua, B.; Lin, J. C.; Williams, V. D.; Kummler, P.; Neidigh, J. W.; Andersen, N. H. The Trp-cage: Optimizing the Stability of a Globular Miniprotein. Protein Eng. Des. Sel. 2008, 21, 171–185.

(82) A. Doyle, D.; Cabral, J. a. M.; Pfuetzner, R. A.; Kuo, A.; Gulbis, J. M.; Cohen, S. L.; Chait, B. T.; MacKinnon, R. The Structure of the Potassium Channel: Molecular Basis of K^+^ Conduction and Selectivity. Science 1998, 280, 69–77.

(83) Tan, L.; Yan, W.; McCorvy, J. D.; Cheng, J. Biased Ligands of G Protein-Coupled Receptors (GPCRs): Structure–Functional Selectivity Relationships (SFSRs) and Therapeutic Potential. J. Med. Chem. 2018, 61, 9841–9878.

(84) Rajagopal, S.; Ahn, S.; Rominger, D. H.; Gowen-MacDonald, W.; Lam, C. M.; DeWire, S. M.; Violin, J. D.; Lefkowitz, R. J. Quantifying Ligand Bias at Seven-Transmembrane Receptors. Mol. Pharmacol. 2011, 80, 367–377.

(85) Strachan, R. T.; Sun, J.-p.; Rominger, D. H.; Violin, J. D.; Ahn, S.; Rojas Bie Thomsen, A.; Zhu, X.; Kleist, A.; Costa, T.; Lefkowitz, R. J. Divergent Transducer-specific Molecular Efficacies Generate Biased Agonism at a G Protein-Coupled Receptor (GPCR). J. Biol. Chem. 2014, 289, 14211–14224.

(86) Suomivuori, C.-M.; Latorraca, N. R.; Wingler, L. M.; Eismann, S.; King, M. C.; Kleinhenz, A. L. W.; Skiba, M. A.; Staus, D. P.; Kruse, A. C.; Lefkowitz, R. J.; Dror, R. O. Molecular Mechanism of Biased Signaling in a Prototypical G Protein–Coupled Receptor. Science 2020, 367, 881–887.

