## Supplementary Material for "Validation of MHPC512: A Publicly Available Special-Purpose Supercomputer for Molecular Dynamics Simulations of Biomolecular Systems"

### Simulation Setups

#### Ensemble Tests

Systematic validation was conducted based on the work by Merz and Shirts.[1] The validated system consisted of 33,333 flexible TIP3P water molecules, with the initial system constructed using packmol.[2] After construction, energy minimization was performed using the steepest descent method, followed by 250 ps of temperature equilibration in an NVT ensemble, with temperature controlled at 300 K using the v-rescale thermostat.[3] Subsequently, 1 ns of volume equilibration was carried out in an NPT ensemble, using the v-rescale and Parrinello-Rahman algorithms to maintain the temperature at 300 K and pressure at 1 atm. Lennard-Jones interactions were truncated at a distance of 1.2 nm. In comparative simulations with GROMACS and MHPC512, electrostatic interactions were calculated using the particle mesh Ewald (PME)[4,5] and $k$-space Gaussian Split Ewald ($k$-GSE) algorithms, respectively, with real-space truncation at 1.2 nm. The time step was set to 1 fs.

#### Dipeptides

The amino acids were described using the AMBER99SB-ILDN force field, and the TIP3P model was chosen for water molecules. Initially, energy minimization was performed using the steepest descent method, followed by 125 ps of temperature equilibration in an NVT ensemble with temperature controlled at 298 K using the Nosé-Hoover chain (NHC) thermostat. Subsequently, 1 ns of volume equilibration was carried out with temperature maintained at 298 K using the v-rescale thermostat and pressure controlled at 1 atm using the Berendsen barostat. The SHAKE algorithm[6] was used to constrain covalent bonds involving hydrogen atoms. These steps were completed using GROMACS. Finally, production simulations were conducted on MHPC512 using an NPT ensemble, with temperature controlled at 298 K using the NHC thermostat and pressure maintained at 1 atm using the Berendsen barostat. The M-SHAKE algorithm was used to constrain covalent bonds involving hydrogen atoms.

Lennard-Jones interactions were truncated at a distance of 1.2 nm, while electrostatic interactions were calculated using the $k$-GSE algorithm with real-space truncation also set to 1.2 nm. The total simulation time for each system was 8 $\mu s$, and the trajectories were saved every 20 ps. Detailed simulation parameters are provided in Table S1.

Table S1: Simulation parameters for dipeptides on MHPC512.

| # of Atoms | System | Ions | Box Size (nm) |
|---|---|---|---|
| 95290 | ACE-A-NME | None | $10.0 \times 10.2 \times 10.1$ |
| 104482 | ACE-R-NME | $Cl^-$ | $10.2 \times 10.2 \times 10.2$ |
| 95257 | ACE-D-NME | $Na^+$ | $9.9 \times 9.9 \times 9.9$ |
| 94366 | ACE-G-NME | None | $9.9 \times 9.9 \times 9.9$ |
| 91829 | ACE-P-NME | None | $9.8 \times 9.8 \times 9.8$ |
| 99597 | ACE-W-NME | None | $10.0 \times 10.0 \times 10.0$ |

For comparison, similar NPT production simulations were conducted using GROMACS, but with much smaller boxes to save computational cost. The leap-frog integrator was used with a total simulation time of 2 $\mu s$ and a time step of 2 fs. Temperature was controlled at 298 K using the v-rescale thermostat, and pressure was maintained at 1 atm using the c-rescale barostat. Long-range electrostatic interactions were calculated using the PME method, with a truncation radius for non-bonded interactions set to 1.0 nm. The SHAKE algorithm was used to constrain all covalent bonds involving hydrogen atoms. The trajectories were saved every 1 ps. Detailed simulation parameters are provided in Table S2.

Table S2: Simulation parameters for dipeptides with GROMACS.

| # of Atoms | System | Ions | Box Size (nm) |
|---|---|---|---|
| 1948 | ACE-A-NME | None | $2.7 \times 2.7 \times 2.7$ |
| 3145 | ACE-R-NME | $Cl^-$ | $3.2 \times 3.2 \times 3.2$ |
| 2326 | ACE-D-NME | $Na^+$ | $2.9 \times 2.9 \times 2.9$ |
| 2251 | ACE-G-NME | None | $2.8 \times 2.8 \times 2.8$ |
| 2060 | ACE-P-NME | None | $2.8 \times 2.8 \times 2.8$ |
| 2766 | ACE-W-NME | None | $3.0 \times 3.0 \times 3.0$ |

#### Folding

##### Chignolin (CLN025)

An extended conformation was generated with AmberTools and parametrized in GROMACS using the AMBER14SB force field. The peptide was solvated in a ∼10 nm cubic TIP3P water box (32,709 water molecules) and neutralized with two $Na^+$ ions, yielding a 98,295-atom system. After energy minimization, the system underwent 500 ps of NVT and 500 ps of NPT equilibration. Subsequently, two 10 $\mu$s NVT production runs were carried out: one on MHPC512 and another in GROMACS for direct comparison. Simulations were maintained at 325 K with an NHC thermostat. Lennard-Jones and short-range Coulomb cutoffs were set to 1.2 nm; long-range electrostatics were treated with the $k$-GSE algorithm on MHPC512 and with PME in GROMACS. A 2.5 fs timestep was used, and coordinates were stored every 400,000 steps.

##### Trp-cage

The linear structure of the sequence DAYAQWLADGGPSSGRPPPS was generated using AmberTools, corresponding to the heat-stable K8A variant TC10b of Trp-cage. The system setup was performed using GROMACS with the CHARMM22∗ force field. The linear peptide was dissolved in a cubic box of TIP3P water with an edge length of 10 nm approximately, containing 32,607 water molecules and 65 mM NaCl, resulting in a total of 98,172 atoms. Subsequently, energy minimization and short NVT and NPT equilibrations (each 500 ps) were performed using GROMACS.

A 20-$\mu$s long NVT production simulation was conducted on MHPC512. During the simulation, the temperature was controlled at 290 K using the NHC thermostat. The cutoff values for Lennard-Jones and short-range electrostatic interactions were set to 1.2 nm, and long-range electrostatic interactions were handled using the $k$-GSE algorithm. The time step was set to 2.5 fs, and trajectory data were saved every 400,000 steps for analysis.

##### Protein B

The linear form of Protein B (sequence LKNAIEDAIAELKKAGITSDFYFNAINKAKTVEEV-NALVNEILKAHA), representing the K5I/K39V double mutant of its binding domain, was built according to Johansson et al.[7] System preparation was carried out in GROMACS by solvating the peptide in a cubic TIP3P water box with an edge length of ∼13 nm. Sodium and chloride ions were added to neutralize total charge and establish a 50 mM NaCl solution, yielding 71,354 water molecules, 67 $Na^+$, and 66 $Cl^-$ (214 932 atoms in total). The peptide was described with the CHARMM22∗ force field. Steepest-descent energy minimization was followed by 100 ps of NVT equilibration and 1 ns of NPT equilibration. The equilibrated system was then propagated for 30 $\mu$s on MHPC512 under NVT conditions. Temperature was held at 340 K with an NHC thermostat. Lennard-Jones and real-space electrostatic interactions were truncated at 1.2 nm, while long-range electrostatics were treated with the $k$-GSE algorithm. A 2.5 fs integration timestep was employed, and frames were saved every 400,000 steps (1 ns).

#### Membrane Systems

Each system was neutralized and adjusted to 0.15 M NaCl. The AMBER system (88,811 atoms) was constructed with packmol-memgen,[8] whereas the CHARMM system (96,020 atoms) was generated with MolCube (https://www.molcube.com/). After steepest-descent energy minimization, the temperature was raised to 300 K over 500 ps with the volume fixed using v-rescale thermostat, followed by 50 ns of NPT equilibration using Berendsen coupling. Post-equilibration box dimensions were 10.18 nm × 10.09 nm × 8.38 nm (AMBER system) and 10.33 nm × 10.33 nm × 8.64 nm (CHARMM system). Identical 200 ns production runs were then initiated on both MHPC512 and GROMACS.

MHPC512 production simulations were carried out in the NPT ensemble with a 2.5 fs velocity-Verlet integrator, a Berendsen thermostat at 300 K, and a semi-isotropic Berendsen barostat at 1 atm. Long-range electrostatics were evaluated with the $k$-GSE method, and van der Waals interactions were truncated at 1.2 nm. All bonds involving hydrogen atoms were constrained with M-SHAKE. Corresponding GROMACS simulations used identical integrator, timestep, temperature, and pressure controls, but employed PME for electrostatics and LINCS[9] for bond constraints.

### Applications

#### Self Assembly of Lipid Molecules in Aqueous Solution

The simulation protocol closely followed that of Skjevik et al.[10] First, 214 molecules of 1,2-dioleoyl-sn-glycero-3-phosphoethanolamine (DOPE) and 72 molecules of 1,2-dioleoyl-sn-glycero-3-phosphoglycerol potassium salt (DOPG) were randomly inserted into a simulation box with the gmx insert-molecules utility in GROMACS. The system was subsequently solvated with 18,750 TIP3P water molecules. Parameterization employed the lipid17 force field and TIP3P water model in AmberTools' tleap, yielding AMBER-format topology and coordinate files that were converted to GROMACS-compatible .top and .gro files with ACPYPE.[11] After 50,000 steps of steepest-descent energy minimization, the system was heated to 310 K over 500 ps under constant volume, followed by 20 ns of isotropic NPT equilibration at 1 bar to relax the density. Production was carried out as a 2 $\mu$s semi-isotropic NPT simulation at 310 K and 1 bar on MHPC512.

#### Conformation Transition between Alternative State and Canonical Active State for $AT_1R$

Building upon previous work,[12] a molecular dynamics simulation was initiated using the crystal structure of the active AT1R–AngII–NbAT110i1 complex (PDB ID: 6OS0[13]) with the nanobody NbAT110i1 removed. The receptor's N- and C-terminal residues were neutralized with acetyl and N-methylamide caps, respectively. Asp74 and Asp125 were protonated;

all other titratable residues were set to their predominant protonation states at pH 7.0. Missing segments were built with SWISS-MODEL.[14] Two disulfide bonds (Cys180–Cys101 and Cys18–Cys274) were retained. The receptor–ligand complex was embedded in a pre-equilibrated POPC bilayer with Membrane Builder (MolCube), oriented according to the OPM database.[15] The system was solvated with TIP3P water, neutralized with 0.15 M NaCl, and parameterized with the CHARMM36m force field. After 5,000 steps of steepest-descent energy minimization, the system underwent 125 ps of NVT equilibration with a Berendsen thermostat at 310 K and positional restraints on the solute. A second 125 ps NVT phase employed the v-rescale thermostat at 310 K with weakened restraints. Four successive NPT equilibration stages (125 ps, 500 ps, 500 ps, 500 ps) were run at 310 K and 1 atm using the v-rescale thermostat and c-rescale barostat, gradually releasing the restraints. All equilibration steps were performed with GROMACS. Production runs (5 $\mu$s) were performed on MHPC512 in the NPT ensemble, using the NHC thermostat (310 K) and a semi-isotropic Parrinello–Rahman barostat (1 atm). Bonds to hydrogens were constrained with SHAKE. Lennard-Jones interactions were cut off at 1.2 nm; electrostatics were treated with the $k$-GSE algorithm with a 1.2 nm real-space cutoff. Trajectories were saved every 20 ps.

### Potential and Virial

#### GROMACS-formatted Input

Table S3: Potential energy and virial (kJ/mol) of a SPC Water Box

| Potential Energy | | | |
|---|---|---|---|
| | GMX | MHPC512 | Relative Error |
| Total | -1366530.00 | -1366515.84 | -0.0010% |
| VDW | 229183.12 | 229182.08 | -0.0005% |
| Ele. | -1595705.23 | -1595697.92 | -0.0005% |
| Virial | | | |
| | GMX | MHPC512 | MHPC512/GMX |
| Total | -2028550.00 | -4057124.16 | 2.0000 |
| x-component | -670485.00 | -1340981.52 | 2.0000 |
| y-component | -679485.00 | -1358979.04 | 2.0000 |
| z-component | -678580.00 | -1357163.52 | 2.0000 |

Table S4: Potential energy and virial (kJ/mol) of a SPC/E Water Box

| Potential Energy | | | |
|---|---|---|---|
| | GMX | MHPC512 | Relative Error |
| Total | -1532820.00 | -1532805.44 | -0.0009% |
| VDW | 290720.07 | 290716.72 | -0.0012% |
| Ele. | -1823530.86 | -1823522.24 | -0.0005% |
| Virial | | | |
| | GMX | MHPC512 | MHPC512/GMX |
| Total | -2361370.00 | -4722762.24 | 2.0000 |
| x-component | -789430.00 | -1578874.40 | 2.0000 |
| y-component | -787742.00 | -1575488.48 | 2.0000 |
| z-component | -784203.00 | -1568399.36 | 2.0000 |

Table S5: Potential energy and virial (kJ/mol) of a CHARMM mTIP3P Water Box

| Potential Energy | | | |
|---|---|---|---|
| | GMX | MHPC512 | Relative Error |
| Total | -1349293.64 | -1349279.51 | -0.0010% |
| VDW | 139726.22 | 139725.09 | -0.0008% |
| Ele. | -1489019.87 | -1489004.60 | -0.0010% |
| Virial | | | |
| | GMX | MHPC512 | MHPC512/GMX |
| Total | -1749294.72 | -3498610.23 | 2.0000 |
| x-component | -582054.26 | -1164122.53 | 2.0000 |
| y-component | -578640.52 | -1157286.49 | 2.0000 |
| z-component | -588599.94 | -1177201.22 | 2.0000 |

Table S6: Potential energy and virial (kJ/mol) of a OPC3 Water Box

| Potential Energy | | | |
|---|---|---|---|
| | GMX | MHPC512 | Relative Error |
| Total | -1636993.94 | -1636985.45 | -0.0005% |
| VDW | 305505.34 | 305505.48 | 0.0000% |
| Ele. | -1942499.28 | -1942490.93 | -0.0004% |
| Virial | | | |
| | GMX | MHPC512 | MHPC512/GMX |
| Total | -2511663.47 | -5023344.70 | 2.0000 |
| x-component | -835064.08 | -1670140.07 | 2.0000 |
| y-component | -836380.65 | -1672756.91 | 2.0000 |
| z-component | -840218.74 | -1680447.72 | 2.0000 |

Table S7: Potential energy and virial (kJ/mol) of JAK2 (PDB ID: 8C09) with CHARMM36 force field

| Potential Energy | | | |
|---|---|---|---|
| | GMX | MHPC512 | Relative Error |
| Total | -1543545.38 | -1543540.96 | -0.0003% |
| VDW | 279079.08 | 279075.76 | -0.0012% |
| Ele. | -1841295.58 | -1841287.84 | -0.0004% |
| Bond+Angle+UB | 13567.61 | 13567.59 | -0.0002% |
| Proper Dih. | 5954.95 | 5954.95 | 0.0000% |
| Improper Dih. | 544.42 | 544.42 | 0.0008% |
| CMAP | -1395.80 | -1395.80 | 0.0001% |
| Virial | | | |
| | GMX | MHPC512 | MHPC512/GMX |
| Total | -2329050.31 | -4658076.80 | 2.0000 |
| x-component | -776970.25 | -1553948.48 | 2.0000 |
| y-component | -773213.31 | -1546405.28 | 2.0000 |
| z-component | -778866.75 | -1557722.88 | 2.0000 |

Table S8: Potential energy and virial (kJ/mol) of JAK2 (PDB ID: 8C09) with CHARMM36m force field

| Potential Energy | | | |
|---|---|---|---|
| | GMX | MHPC512 | Relative Error |
| Total | -1536217.00 | -1536214.08 | -0.0002% |
| VDW | 280332.21 | 280330.32 | -0.0007% |
| Ele. | -1841745.84 | -1841740.96 | -0.0003% |
| Bond+Angle+UB | 13739.30 | 13739.31 | 0.0001% |
| Proper Dih. | 11258.84 | 11258.84 | 0.0000% |
| Improper Dih. | 593.94 | 593.95 | 0.0009% |
| CMAP | -395.51 | -395.51 | -0.0002% |
| Virial | | | |
| | GMX | MHPC512 | MHPC512/GMX |
| Total | -2336183.88 | -4672351.04 | 2.0000 |
| x-component | -781190.38 | -1562371.20 | 2.0000 |
| y-component | -776775.19 | -1553545.44 | 2.0000 |
| z-component | -778218.31 | -1556434.40 | 2.0000 |

Table S9: Potential energy and virial (kJ/mol) of JAK2 (PDB ID: 8C09) with CHARMM27 force field

| Potential Energy | | | |
|---|---|---|---|
| | GMX | MHPC512 | Relative Error |
| Total | -1542400.75 | -1542393.60 | -0.0005% |
| VDW | 279408.39 | 279405.36 | -0.0011% |
| Ele. | -1840644.39 | -1840634.08 | -0.0006% |
| Bond+Angle+UB | 13658.23 | 13658.24 | 0.0000% |
| Proper Dih. | 5991.61 | 5991.62 | 0.0001% |
| Improper Dih. | 578.47 | 578.47 | -0.0003% |
| CMAP | -1393.11 | -1393.12 | 0.0004% |
| Virial | | | |
| | GMX | MHPC512 | MHPC512/GMX |
| Total | -2326795.25 | -4653534.08 | 2.0000 |
| x-component | -773681.00 | -1547355.20 | 2.0000 |
| y-component | -776565.19 | -1553115.04 | 2.0000 |
| z-component | -776549.06 | -1553064.00 | 2.0000 |

Table S10: Potential energy and virial (kJ/mol) of villin (PDB ID: 2F4K) with CHARMM22* force field

| Potential Energy | | | |
|---|---|---|---|
| | GMX | MHPC512 | Relative Error |
| Total | -1384952.88 | -1384935.04 | -0.0013% |
| VDW | 186067.29 | 186064.82 | -0.0013% |
| Ele. | -1573503.91 | -1573483.52 | -0.0013% |
| Bond+Angle+UB | 2122.29 | 2122.27 | -0.0007% |
| Proper Dih. | 361.68 | 361.68 | -0.0012% |
| Improper Dih. | 63.13 | 63.13 | 0.0021% |
| CMAP | -63.39 | -63.39 | 0.0046% |
| Virial | | | |
| | GMX | MHPC512 | MHPC512/GMX |
| Total | -1750391.88 | -3500723.84 | 2.0000 |
| x-component | -579761.50 | -1159510.32 | 2.0000 |
| y-component | -586057.69 | -1172090.24 | 2.0000 |
| z-component | -584572.69 | -1169123.36 | 2.0000 |

Table S11: Potential energy and virial (kJ/mol) of JAK2 (PDB ID: 8C09) with AMBER99SB-ILDN force field

| Potential Energy | | | |
|---|---|---|---|
| | GMX | MHPC512 | Relative Error |
| Total | -1544746.72 | -1544730.88 | -0.0010% |
| VDW | 281112.41 | 281111.36 | -0.0004% |
| Ele. | -1850649.92 | -1850632.96 | -0.0009% |
| Bond | 3577.30 | 3577.31 | 0.0003% |
| Angle | 9531.25 | 9531.25 | 0.0000% |
| Dihedral | 11682.24 | 11682.24 | 0.0000% |
| Virial | | | |
| | GMX | MHPC512 | MHPC512/GMX |
| Total | -2344792.56 | -4689594.56 | 2.0000 |
| x-component | -790014.75 | -1580040.32 | 2.0000 |
| y-component | -776311.06 | -1552633.76 | 2.0000 |
| z-component | -778466.75 | -1556920.48 | 2.0000 |

Table S12: Potential energy and virial (kJ/mol) of JAK2 (PDB ID: 8C09) with AMBER14SB force field

| Potential Energy | | | |
|---|---|---|---|
| | GMX | MHPC512 | Relative Error |
| Total | -1542680.38 | -1542671.52 | -0.0006% |
| VDW | 276450.62 | 276448.92 | -0.0006% |
| Ele. | -1846737.46 | -1846726.88 | -0.0006% |
| Bond | 3431.63 | 3431.62 | -0.0002% |
| Angle | 9412.09 | 9412.09 | 0.0000% |
| Dihedral | 14762.83 | 14762.84 | 0.0000% |
| Virial | | | |
| | GMX | MHPC512 | MHPC512/GMX |
| Total | -2316151.63 | -4632275.84 | 2.0000 |
| x-component | -767982.06 | -1535954.72 | 2.0000 |
| y-component | -778379.75 | -1556757.60 | 2.0000 |
| z-component | -769789.81 | -1539563.52 | 2.0000 |

Table S13: Potential energy and virial (kJ/mol) of JAK2 (PDB ID: 8C09) with AMBER19SB force field

| Potential Energy | | | |
|---|---|---|---|
| | GMX | MHPC512 | Relative Error |
| Total | -1453964.25 | -1453948.64 | -0.0011% |
| VDW | 196528.60 | 196526.56 | -0.0010% |
| Ele. | -1670932.68 | -1670915.04 | -0.0011% |
| Bond | 3479.88 | 3479.87 | -0.0002% |
| Angle | 9814.15 | 9814.15 | -0.0000% |
| Dihedral | 6269.77 | 6269.78 | 0.0002% |
| CMAP | 875.96 | 875.95 | -0.0009% |
| Virial | | | |
| | GMX | MHPC512 | MHPC512/GMX |
| Total | -1855130.19 | -3710235.52 | 2.0000 |
| x-component | -617889.63 | -1235772.96 | 2.0000 |
| y-component | -622720.81 | -1245418.40 | 2.0000 |
| z-component | -614519.75 | -1229044.16 | 2.0000 |

Table S14: Potential energy and virial (kJ/mol) of T4 Lysozyme/ligand complex (PDB ID: 3HTB) with AMBER14SB+GAFF2 force field

| Potential Energy | | | |
|---|---|---|---|
| | GMX | MHPC512 | Relative Error |
| Total | -1426205.63 | -1426190.72 | -0.0010% |
| VDW | 206813.10 | 206812.02 | -0.0005% |
| Ele. | -1649267.33 | -1649251.20 | -0.0010% |
| Bond | 2112.77 | 2112.76 | -0.0005% |
| Angle | 5356.36 | 5356.36 | 0.0000% |
| Dihedral | 8779.46 | 8779.46 | 0.0000% |
| Virial | | | |
| | GMX | MHPC512 | MHPC512/GMX |
| Total | -1954716.69 | -3909446.72 | 2.0000 |
| x-component | -651638.31 | -1303272.48 | 2.0000 |
| y-component | -656153.38 | -1312300.64 | 2.0000 |
| z-component | -646925.00 | -1293873.68 | 2.0000 |

Table S15: Potential energy and virial (kJ/mol) of POPC bilayer built by packmol-memgen with Lipid17 force field

| Potential Energy | | | |
|---|---|---|---|
| | GMX | MHPC512 | Relative Error |
| Total | -1116887.75 | -1116871.52 | -0.0015% |
| VDW | 69937.30 | 69937.60 | 0.0004% |
| Ele. | -1340763.25 | -1340747.36 | -0.0012% |
| Bond | 20909.45 | 20909.43 | -0.0001% |
| Angle | 84225.70 | 84225.77 | 0.0001% |
| Dihedral | 48803.03 | 48803.06 | 0.0001% |
| Virial | | | |
| | GMX | MHPC512 | MHPC512/GMX |
| Total | -1066057.09 | -2132048.16 | 1.9999 |
| x-component | -364265.00 | -728502.40 | 1.9999 |
| y-component | -361560.75 | -723102.56 | 1.9999 |
| z-component | -340231.34 | -680443.12 | 1.9999 |

Table S16: Potential energy and virial (kJ/mol) of POPC bilayer built by packmol-memgen with Lipid21 force field

| Potential Energy | | | |
|---|---|---|---|
| | GMX | MHPC512 | Relative Error |
| Total | -1229484.50 | -1229473.12 | -0.0009% |
| VDW | 61950.63 | 61950.44 | -0.0003% |
| Ele. | -1480797.03 | -1480785.44 | -0.0008% |
| Bond | 24800.22 | 24800.23 | 0.0000% |
| Angle | 103392.46 | 103392.56 | 0.0001% |
| Dihedral | 61169.12 | 61169.15 | 0.0001% |
| Virial | | | |
| | GMX | MHPC512 | MHPC512/GMX |
| Total | -1163739.28 | -2327443.20 | 2.0000 |
| x-component | -394518.06 | -789021.76 | 2.0000 |
| y-component | -390177.31 | -780335.76 | 2.0000 |
| z-component | -379043.91 | -758085.68 | 2.0000 |

Table S17: Potential energy and virial (kJ/mol) of JAK2 (PDB ID: 8C09) with OPLS/AA force field

| Potential Energy | | | |
|---|---|---|---|
| | GMX | MHPC512 | Relative Error |
| Total | -1565547.50 | -1565534.72 | -0.0008% |
| VDW | 282901.42 | 282900.16 | -0.0004% |
| Ele. | -1865186.12 | -1865172.16 | -0.0007% |
| Bond | 3267.31 | 3267.30 | -0.0002% |
| Angle | 8804.78 | 8804.77 | -0.0001% |
| Proper Dih. | 475.48 | 475.48 | 0.0004% |
| Ryckaert-Bell | 4189.65 | 4189.66 | 0.0003% |
| Virial | | | |
| | GMX | MHPC512 | MHPC512/GMX |
| Total | -2348476.31 | -4696960.00 | 2.0000 |
| x-component | -779298.31 | -1558612.16 | 2.0000 |
| y-component | -783817.50 | -1567636.48 | 2.0000 |
| z-component | -785360.50 | -1570711.36 | 2.0000 |

Table S18: Potential energy and virial (kJ/mol) of DNA solution (PDB ID: 1BNA) with DNA.OL21 force field

| Potential Energy | | | |
|---|---|---|---|
| | GMX | MHPC512 | Relative Error |
| Total | -1303358.63 | -1303349.76 | -0.0007% |
| VDW | 188639.94 | 188639.44 | -0.0003% |
| Ele. | -1497472.66 | -1497463.36 | -0.0006% |
| Bond | 756.66 | 756.66 | 0.0003% |
| Angle | 1846.95 | 1846.95 | 0.0001% |
| Dihedral | 2870.49 | 2870.49 | -0.0001% |
| Virial | | | |
| | GMX | MHPC512 | MHPC512/GMX |
| Total | -1767882.63 | -3535801.60 | 2.0000 |
| x-component | -589576.56 | -1179150.32 | 2.0000 |
| y-component | -588542.81 | -1177110.80 | 2.0000 |
| z-component | -589763.25 | -1179540.64 | 2.0000 |

Table S19: Potential energy and virial (kJ/mol) of DNA solution (PDB ID: 1BNA) with DNA.OL24 force field

| Potential Energy | | | |
|---|---|---|---|
| | GMX | MHPC512 | Relative Error |
| Total | -1300116.00 | -1300104.00 | -0.0009% |
| VDW | 189557.60 | 189555.88 | -0.0009% |
| Ele. | -1498578.10 | -1498564.32 | -0.0009% |
| Bond | 879.17 | 879.18 | 0.0009% |
| Angle | 1641.89 | 1641.89 | -0.0003% |
| Dihedral | 6383.48 | 6383.48 | 0.0000% |
| Virial | | | |
| | GMX | MHPC512 | MHPC512/GMX |
| Total | -1772842.50 | -3545656.96 | 2.0000 |
| x-component | -590593.06 | -1181185.68 | 2.0000 |
| y-component | -591619.31 | -1183224.72 | 2.0000 |
| z-component | -590630.13 | -1181246.40 | 2.0000 |

Table S20: Potential energy and virial (kJ/mol) of RNA solution (PDB ID: 1ESH) with DNA.OL3 force field

| Potential Energy | | | |
|---|---|---|---|
| | GMX | MHPC512 | Relative Error |
| Total | -1388562.63 | -1388553.28 | -0.0007% |
| VDW | 203625.55 | 203624.24 | -0.0006% |
| Ele. | -1595000.55 | -1594989.92 | -0.0007% |
| Bond | 471.54 | 471.53 | -0.0014% |
| Angle | 995.34 | 995.34 | 0.0000% |
| Dihedral | 1345.46 | 1345.45 | -0.0004% |
| Virial | | | |
| | GMX | MHPC512 | MHPC512/GMX |
| Total | -1900773.88 | -3801508.48 | 2.0000 |
| x-component | -639548.94 | -1279070.16 | 2.0000 |
| y-component | -629694.25 | -1259382.56 | 2.0000 |
| z-component | -631530.69 | -1263055.92 | 2.0000 |

Table S21: Potential energy and virial (kJ/mol) of glycoprotein (PDB ID: 3CFW) with GLYCAM_06j-1 force field built on MolCube

| Potential Energy | | | |
|---|---|---|---|
| | GMX | MHPC512 | Relative Error |
| Total | -1451598.50 | -1451601.44 | 0.0002% |
| VDW | 204760.25 | 204753.48 | -0.0033% |
| Ele. | -1672212.77 | -1672208.96 | -0.0002% |
| Bond | 2219.02 | 2219.01 | -0.0007% |
| Angle | 5273.87 | 5273.87 | 0.0000% |
| Dihedral | 8361.06 | 8361.07 | 0.0001% |
| Virial | | | |
| | GMX | MHPC512 | MHPC512/GMX |
| Total | -1897166.94 | -3794185.60 | 1.9999 |
| x-component | -632422.06 | -1264796.96 | 1.9999 |
| y-component | -633647.38 | -1267243.60 | 1.9999 |
| z-component | -631097.50 | -1262144.80 | 1.9999 |

#### AMBER-formatted Input

Table S22: Potential energy (kJ/mol) of a SPC Water Box

| | AMBER | MHPC512 | Relative Error |
|---|---|---|---|
| Total | -1363564.80 | -1363607.44 | -0.0031% |
| VDW | 227163.60 | 227163.46 | 0.0001% |
| Ele. | -1590728.48 | -1590766.43 | -0.0024% |

Table S23: Potential energy (kJ/mol) of a SPC/E Water Box

| | AMBER | MHPC512 | Relative Error |
|---|---|---|---|
| Total | -1533603.36 | -1533540.00 | -0.0041% |
| VDW | 290066.96 | 290067.10 | 0.0000% |
| Ele. | -1823656.50 | -1823607.20 | -0.0027% |

Table S24: Potential energy (kJ/mol) of a TIP3P Water Box

| | AMBER | MHPC512 | Relative Error |
|---|---|---|---|
| Total | -1312604.48 | -1312580.08 | -0.0019% |
| VDW | 196408.76 | 196408.94 | 0.0001% |
| Ele. | -1509022.66 | -1508989.12 | -0.0022% |

Table S25: Potential energy (kJ/mol) of JNK3 (PDB ID: 4W4W) with AMBER14SB force field

| | AMBER | MHPC512 | Relative Error |
|---|---|---|---|
| Total | -1099336.16 | -1099307.60 | -0.0026% |
| Bond | 4603.32 | 4603.33 | 0.0002% |
| Angle | 12391.34 | 12391.34 | 0.0000% |
| Dihedral | 19449.69 | 19449.70 | 0.0001% |
| VDW | 142474.09 | 147956.75 | 0.0001% |
| 1-4 VDW | 5482.56 | | |
| Ele. | -1344270.98 | -1283708.72 | -0.0022% |
| 1-4 Ele. | 60533.83 | | |

Table S26: Potential energy (kJ/mol) of JNK3 (PDB ID: 4W4W) with AMBER19SB force field

| | AMBER | MHPC512 | Relative Error |
|---|---|---|---|
| Total | -1110703.43 | -1110671.52 | -0.0029% |
| Bond | 4489.67 | 4489.68 | 0.0003% |
| Angle | 12220.91 | 12220.90 | 0.0000% |
| Dihedral | 8422.18 | 8422.19 | 0.0001% |
| VDW | 143422.11 | 148886.75 | 0.0001% |
| 1-4 VDW | 5464.54 | | |
| Ele. | -1346397.08 | -1285939.76 | -0.0025% |
| 1-4 Ele. | 60425.51 | | |
| CMAP | 1248.76 | 1248.75 | -0.0004% |

Table S27: Potential energy (kJ/mol) of T4 Lysozyme/ligand complex (PDB ID: 3HTB) with AMBER14SB+GAFF2 force field

<table>
<tr><th></th><th>AMBER</th><th>MHPC512</th><th>Relative Error</th></tr>
<tr><td>Total</td><td>-1333064.86</td><td>-1333042.56</td><td>-0.0017%</td></tr>
<tr><td>Bond</td><td>2120.41</td><td>2120.41</td><td>0.0000%</td></tr>
<tr><td>Angle</td><td>5447.08</td><td>5447.08</td><td>-0.0001%</td></tr>
<tr><td>Dihedral</td><td>8881.86</td><td>8881.86</td><td>0.0001%</td></tr>
<tr><td>VDW</td><td>190508.50</td><td rowspan="2">193022.36</td><td rowspan="2">0.0001%</td></tr>
<tr><td>1-4 VDW</td><td>2513.63</td></tr>
<tr><td>Ele.</td><td>-1564690.40</td><td rowspan="2">-1542514.24</td><td rowspan="2">-0.0014%</td></tr>
<tr><td>1-4 Ele.</td><td>22154.06</td></tr>
</table>

Table S28: Potential energy (kJ/mol) of POPC bilayer with Lipid17 force field built by packmol-memgen

<table>
<tr><th></th><th>AMBER</th><th>MHPC512</th><th>Relative Error</th></tr>
<tr><td>Total</td><td>-1051764.32</td><td>-1051723.36</td><td>-0.0039%</td></tr>
<tr><td>Bond</td><td>21472.59</td><td>21472.57</td><td>-0.0001%</td></tr>
<tr><td>Angle</td><td>88044.38</td><td>88044.40</td><td>0.0000%</td></tr>
<tr><td>Dihedral</td><td>52175.46</td><td>52175.48</td><td>0.0000%</td></tr>
<tr><td>VDW</td><td>60270.96</td><td rowspan="2">78145.71</td><td rowspan="2">0.0002%</td></tr>
<tr><td>1-4 VDW</td><td>17874.59</td></tr>
<tr><td>Ele.</td><td>-885556.02</td><td rowspan="2">-1291561.52</td><td rowspan="2">-0.0032%</td></tr>
<tr><td>1-4 Ele.</td><td>-406046.28</td></tr>
</table>

Table S29: Potential energy (kJ/mol) of POPC bilayer with Lipid21 force field built on MolCube

<table>
<tr><th></th><th>AMBER</th><th>MHPC512</th><th>Relative Error</th></tr>
<tr><td>Total</td><td>-1121354.95</td><td>-1121322.88</td><td>-0.0029%</td></tr>
<tr><td>Bond</td><td>24419.92</td><td>24419.89</td><td>-0.0001%</td></tr>
<tr><td>Angle</td><td>97235.56</td><td>97235.64</td><td>0.0001%</td></tr>
<tr><td>Dihedral</td><td>60028.33</td><td>60028.35</td><td>0.0000%</td></tr>
<tr><td>VDW</td><td>50695.08</td><td rowspan="2">71714.18</td><td rowspan="2">0.0006%</td></tr>
<tr><td>1-4 VDW</td><td>21018.67</td></tr>
<tr><td>Ele.</td><td>-1207356.47</td><td rowspan="2">-1375165.92</td><td rowspan="2">-0.0023%</td></tr>
<tr><td>1-4 Ele.</td><td>-167840.99</td></tr>
<tr><td>CMAP</td><td>444.95</td><td>444.95</td><td>-0.0009%</td></tr>
</table>

Table S30: Potential energy (kJ/mol) of DNA solution (PDB ID: 1BNA) with Parmbsc1 force field

| | AMBER | MHPC512 | Relative Error |
|---|---:|---:|---:|
| Total | -1346388.62 | -1346352.01 | -0.0027% |
| Bond | 893.30 | 893.30 | 0.0005% |
| Angle | 1807.84 | 1807.83 | -0.0004% |
| Dihedral | 2422.33 | 2422.33 | 0.0000% |
| VDW | 198891.33 | 198891.11 | -0.0001% |
| Ele. | -1550403.42 | -1550366.58 | -0.0024% |

Table S31: Potential energy (kJ/mol) of DNA solution (PDB ID: 1BNA) with DNA.OL15 force field

| | AMBER | MHPC512 | Relative Error |
|---|---:|---:|---:|
| Total | -1347462.22 | -1347424.94 | -0.0028% |
| Bond | 841.18 | 841.18 | 0.0003% |
| Angle | 1871.81 | 1871.80 | -0.0003% |
| Dihedral | 2845.14 | 2845.14 | 0.0000% |
| VDW | 198386.90 | 198386.64 | -0.0001% |
| Ele. | -1551407.24 | -1551369.70 | -0.0024% |

Table S32: Potential energy (kJ/mol) of DNA solution (PDB ID: 1BNA) with DNA.OL21 force field

| | AMBER | MHPC512 | Relative Error |
|---|---:|---:|---:|
| Total | -1349510.55 | -1349467.20 | -0.0032% |
| Bond | 695.41 | 695.41 | -0.0006% |
| Angle | 1711.91 | 1711.91 | 0.0000% |
| Dihedral | 3006.40 | 3006.40 | 0.0001% |
| VDW | 195572.33 | 196398.64 | 0.0000% |
| 1-4 VDW | 826.22 | | |
| Ele. | -1538014.59 | -1551280.16 | -0.0028% |
| 1-4 Ele. | -13308.23 | | |

Table S33: Potential energy (kJ/mol) of DNA solution (PDB ID: 1BNA) with DNA.OL24 force field

<table>
<tr><th></th><th>AMBER</th><th>MHPC512</th><th>Relative Error</th></tr>
<tr><td>Total</td><td>-1351162.21</td><td>-1351119.20</td><td>-0.0032%</td></tr>
<tr><td>Bond</td><td>757.06</td><td>757.05</td><td>-0.0011%</td></tr>
<tr><td>Angle</td><td>1758.36</td><td>1758.37</td><td>0.0003%</td></tr>
<tr><td>Dihedral</td><td>6376.21</td><td>6376.21</td><td>0.0000%</td></tr>
<tr><td>VDW</td><td>197594.72</td><td rowspan="2">198466.48</td><td rowspan="2">0.0000%</td></tr>
<tr><td>1-4 VDW</td><td>871.70</td></tr>
<tr><td>Ele.</td><td>-1545237.02</td><td rowspan="2">-1558477.40</td><td rowspan="2">-0.0027%</td></tr>
<tr><td>1-4 Ele.</td><td>-13283.23</td></tr>
</table>

Table S34: Potential energy (kJ/mol) of RNA solution (PDB ID: 1ESH) with RNA.OL3 force field

<table>
<tr><th></th><th>AMBER</th><th>MHPC512</th><th>Relative Error</th></tr>
<tr><td>Total</td><td>-1363076.65</td><td>-1363038.08</td><td>-0.0028%</td></tr>
<tr><td>Bond</td><td>435.56</td><td>435.56</td><td>-0.0003%</td></tr>
<tr><td>Angle</td><td>958.66</td><td>958.66</td><td>-0.0001%</td></tr>
<tr><td>Dihedral</td><td>1375.23</td><td>1375.23</td><td>0.0002%</td></tr>
<tr><td>VDW</td><td>202083.51</td><td rowspan="2">202591.20</td><td rowspan="2">0.0001%</td></tr>
<tr><td>1-4 VDW</td><td>507.59</td></tr>
<tr><td>Ele.</td><td>-1562088.97</td><td rowspan="2">-1568398.72</td><td rowspan="2">-0.0025%</td></tr>
<tr><td>1-4 Ele.</td><td>-6348.22</td></tr>
</table>

Table S35: Potential energy (kJ/mol) of RNA solution (PDB ID: 1ESH) with RNA.Shaw force field

| | AMBER | MHPC512 | Relative Error |
|---|---|---|---|
| Total | -1344451.29 | -1344413.32 | -0.0028% |
| Bond | 480.47 | 480.47 | 0.0009% |
| Angle | 830.39 | 830.39 | 0.0004% |
| Dihedral | 1173.73 | 1173.73 | -0.0001% |
| VDW | 202574.97 | 202574.70 | -0.0001% |
| Ele. | -1549510.84 | -1549472.61 | -0.0025% |

Table S36: Potential energy (kJ/mol) of JAK2 (PDB ID: 8C09) with CHARMM36m force field

| | AMBER | MHPC512 | Relative Error |
|---|---|---|---|
| Total | -1344306.53 | -1344272.48 | -0.0025% |
| Bond | 3642.44 | 4849.16 | 0.0002% |
| U-B | 1206.71 | | |
| Angle | 9222.57 | 9222.57 | 0.0000% |
| Dihedral | 11659.83 | 11659.84 | 0.0001% |
| CMAP | -436.58 | -436.58 | 0.0002% |
| Improper Dihedral | 585.88 | 585.88 | 0.0005% |
| VDW | 120570.06 | 123795.80 | -0.0008% |
| 1-4 VDW | 3226.73 | | |
| Ele. | -1541426.45 | -1493949.12 | -0.0023% |
| 1-4 Ele. | 47442.29 | | |

Table S37: Potential energy (kJ/mol) of oligosaccharide with GLYCAM_06j-1 force field built on MolCube

| | AMBER | MHPC512 | Relative Error |
|---|---|---|---|
| Total | -1259207.91 | -1259182.64 | -0.0020% |
| Bond | 276.51 | 276.51 | -0.0006% |
| Angle | 624.50 | 624.50 | 0.0002% |
| Dihedral | 163.15 | 163.15 | 0.0000% |
| VDW | 189732.33 | 190059.60 | -0.0021% |
| 1-4 VDW | 331.33 | | |
| Ele. | -1459023.11 | -1450306.40 | -0.0020% |
| 1-4 Ele. | 8687.38 | | |

Table S38: Potential energy (kJ/mol) of JNK3 (PDB ID: 4W4W) with phosphorylated serine with phosaa10 force field

| | AMBER | MHPC512 | Relative Error |
|---|---|---|---|
| Total | -1125548.27 | -1125512.80 | 0.0032% |
| Bond | 4847.50 | 4847.51 | 0.0002% |
| Angle | 12972.07 | 12972.07 | 0.0000% |
| Dihedral | 17206.34 | 17206.35 | 0.0000% |
| VDW | 143416.43 | 148960.96 | 0.0001% |
| 1-4 VDW | 5544.43 | | |
| Ele. | -1365607.89 | -1309499.68 | 0.0027% |
| 1-4 Ele. | 56072.86 | | |

Table S39: Potential energy (kJ/mol) of JNK3 (PDB ID: 4W4W) with phosphorylated serine with phosaa14 force field

| | AMBER | MHPC512 | Relative Error |
|---|---|---|---|
| Total | -1120235.72 | -1120204.40 | 0.0028% |
| Bond | 4719.82 | 4719.81 | 0.0001% |
| Angle | 13185.40 | 13185.41 | 0.0001% |
| Dihedral | 21192.43 | 21192.44 | 0.0000% |
| VDW | 143429.29 | 149021.24 | 0.0001% |
| 1-4 VDW | 5591.85 | | |
| Ele. | -1364778.69 | -1308323.36 | 0.0024% |
| 1-4 Ele. | 56424.19 | | |

Table S40: Potential energy (kJ/mol) of JNK3 (PDB ID: 4W4W) with phosphorylated serine with phosaa19 force field

| | AMBER | MHPC512 | Relative Error |
|---|---|---|---|
| Total | -1130128.35 | -1130096.80 | 0.0028% |
| Bond | 4609.75 | 4609.76 | 0.0002% |
| Angle | 13047.18 | 13047.17 | 0.0001% |
| Dihedral | 10290.65 | 10290.66 | 0.0001% |
| VDW | 144947.25 | 150440.14 | 0.0001% |
| 1-4 VDW | 5492.79 | | |
| Ele. | -1366128.96 | -1309693.68 | 0.0024% |
| 1-4 Ele. | 56403.87 | | |
| CMAP | 1209.13 | 1209.13 | 0.0000% |

Table S41: Potential energy (kJ/mol) of DNA solution (PDB ID: 1BNA) with Tumuc1 DNA force field

| | AMBER | MHPC512 | Relative Error |
|---|---|---|---|
| Total | -1409765.27 | -1409727.57 | -0.0027% |
| Bond | 1044.97 | 1044.98 | 0.0010% |
| Angle | 1860.42 | 1860.42 | 0.0000% |
| Dihedral | 2824.69 | 2824.69 | 0.0001% |
| VDW | 857.51 | 206993.05 | -0.0002% |
| 1-4 VDW | 206135.85 | | |
| Ele. | -27946.28 | -1622488.72 | -0.0023% |
| 1-4 Ele. | -1594542.40 | | |

### References


(1) Merz, P. T.; Shirts, M. R. Testing for Physical Validity in Molecular Simulations. *PLOS One* **2018**, *13*, 1–22.

(2) Martínez, L.; Andrade, R.; Birgin, E. G.; Martínez, J. M. PACKMOL: A Package for Building Initial Configurations for Molecular Dynamics Simulations. *J. Comput. Chem.* **2009**, *30*, 2157–2164.

(3) Bussi, G.; Donadio, D.; Parrinello, M. Canonical Sampling through Velocity Rescaling. *J. Chem. Phys.* **2007**, *126*, 014101.

(4) Darden, T.; York, D.; Pedersen, L. Particle Mesh Ewald: An $N \cdot \log(N)$ Method for Ewald Sums in Large Systems. *J. Chem. Phys.* **1993**, *98*, 10089–10092.

(5) Essmann, U.; Perera, L.; Berkowitz, M. L.; Darden, T.; Lee, H.; Pedersen, L. G. A Smooth Particle Mesh Ewald Method. *J. Chem. Phys.* **1995**, *103*, 8577–8593.

(6) Ryckaert, J.-P.; Ciccotti, G.; Berendsen, H. J. Numerical Integration of the Cartesian Equations of Motion of a System with Constraints: Molecular Dynamics of n-Alkanes. *J. Comput. Phys.* **1977**, *23*, 327–341.

(7) Johansson, M. U.; de Château, M.; Wikström, M.; Forsén, S.; Drakenberg, T.; Björck, L. Solution Structure of the Albumin-binding GA Module: A Versatile Bacterial Protein Domain. *J. Mol. Biol.* **1997**, *266*, 859–865.

(8) Schott-Verdugo, S.; Gohlke, H. PACKMOL-Memgen: A Simple-To-Use, Generalized Workflow for Membrane-Protein–Lipid-Bilayer System Building. *J. Chem. Inf. Model.* **2019**, *59*, 2522–2528.

(9) Hess, B.; Bekker, H.; Berendsen, H. J. C.; Fraaije, J. G. E. M. LINCS: A Linear Constraint Solver for Molecular Simulations. *J. Comput. Chem.* **1997**, *18*, 1463–1472.

(10) Skjevik, Å. A.; Madej, B. D.; Dickson, C. J.; Teigen, K.; Walker, R. C.; Gould, I. R. All-atom Lipid Bilayer Self-Assembly with the AMBER and CHARMM Lipid Force Fields. *Chem. Commun.* **2015**, *51*, 4402–4405.

(11) Sousa da Silva, A. W.; Vranken, W. F. ACPYPE-Antechamber Python Parser Interface. *BMC Res. Notes* **2012**, *5*, 1–8.

(12) Suomivuori, C.-M.; Latorraca, N. R.; Wingler, L. M.; Eismann, S.; King, M. C.; Kleinhenz, A. L. W.; Skiba, M. A.; Staus, D. P.; Kruse, A. C.; Lefkowitz, R. J.; Dror, R. O. Molecular Mechanism of Biased Signaling in a Prototypical G Protein–Coupled Receptor. *Science* **2020**, *367*, 881–887.

(13) Wingler, L. M.; Skiba, M. A.; McMahon, C.; Staus, D. P.; Kleinhenz, A. L. W.; Suomivuori, C.-M.; Latorraca, N. R.; Dror, R. O.; Lefkowitz, R. J.; Kruse, A. C. Angiotensin and Biased Analogs Induce Structurally Distinct Active Conformations within a GPCR. *Science* **2020**, *367*, 888–892.

(14) Schwede, T.; Kopp, J.; Guex, N.; Peitsch, M. C. SWISS-MODEL: An Automated Protein Homology-Modeling Server. *Nucleic Acids Res.* **2003**, *31*, 3381–3385.

(15) Lomize, M. A.; Pogozheva, I. D.; Joo, H.; Mosberg, H. I.; Lomize, A. L. OPM Database and PPM Web Server: Resources for Positioning of Proteins in Membranes. *Nucleic Acids Res.* **2011**, *40*, D370–D376.
